# Consequences of endogenous and virally-induced hyperphosphorylated tau on behavior and cognition in a rat model of Alzheimer’s disease

**DOI:** 10.1101/2021.10.30.466530

**Authors:** Michael A. Kelberman, Claire R. Anderson, Eli Chlan, Jacki M. Rorabaugh, Katharine E. McCann, David Weinshenker

## Abstract

**Background:** The locus coeruleus (LC) is one of the earliest brain regions to accumulate hyperphosphorylated tau, but a lack of animal models that recapitulate this pathology has hampered our ability to understand its contributions to the pathophysiology of Alzheimer’s disease (AD).

**Objective:** We previously reported that TgF344-AD rats, which overexpress mutant human amyloid precursor protein and presenilin-1, accumulate early endogenous hyperphosphorylated tau in the LC. Here, we used TgF344-AD rats and a wild-type (WT) human tau virus to interrogate the effects of endogenous hyperphosphorylated rat tau and human tau in the LC on AD-related neuropathology and behavior.

**Methods:** Two-month old TgF344-AD and WT rats received bilateral LC infusions of WT human tau or mCherry control virus driven by the noradrenergic-specific PRSx8 promoter. Rats were subsequently assessed at 6 and 12 months for arousal (sleep latency), anxiety-like behavior (open field, elevated plus maze, novelty-suppressed feeding), passive coping (forced swim task), and learning and memory (Morris water maze and fear conditioning). Hippocampal microglia, astrocyte, and AD pathology were evaluated using immunohistochemistry.

**Results:** In general, the effects of age were more pronounced than genotype or treatment; older rats displayed greater hippocampal pathology, took longer to fall asleep, had reduced locomotor activity, floated more, and had impaired cognition compared to younger animals. TgF344-AD rats showed increased anxiety-like behavior and impaired learning and memory. The tau virus had negligible influence on most measures.

**Conclusion:** Effects of hyperphosphorylated tau on AD-like neuropathology and behavioral symptoms were subtle. Further investigation of different forms of tau is warranted.

## Introduction

Alzheimer’s disease (AD) is the most common neurodegenerative disorder in the world and is characterized by two protein aggregates: amyloid-β (Aβ) plaques and neurofibrillary tangles composed of hyperphosphorylated and misfolded tau. Initiation of AD has been attributed to Aβ [1], which subsequently leads to tau hyperphosphorylation [2–4], oxidative stress and mitochondrial dysfunction [5–9], and synaptic impairments [10–12]. However, antibody treatments targeting late-stage Aβ deposition have been largely unsuccessful and/or controversial, highlighting the pressing need to investigate other therapeutic avenues that focus on earlier stages of disease prior to catastrophic neuronal loss and significant cognitive impairment.

Neurofibrillary tangles (NFT) have garnered increasing attention in recent years because tau load is consistently reported as a better predictor of cognitive decline and neuronal degeneration compared with Aβ [13–15]. Although Aβ plaques appear prior to NFT, recent evidence places the deposition of “pretangle” hyperphosphorylated tau prior to both pathologies. Specifically, Braak and colleagues identified the locus coeruleus (LC) as the first brain region to accumulate hyperphosphorylated tau [16], a finding that has since been independently confirmed by other groups [17, 18]. The LC is the primary source of norepinephrine (NE) in the brain, which has potent anti-inflammatory/neuroprotective properties and regulates attention, arousal, and stress responses [19–22], all of which often go awry in AD [23–29]. Although tau pathology can cause cell death and there is evidence for some early reduction of LC volume (which likely corresponds to loss of proximal axons/dendrites) [18, 30–32], frank LC cell body degeneration is not evident until mid- to late-stages of AD [18, 33, 34]. Thus, detrimental effects of hyperphosphorylated tau pathology on LC function could persist for years or even decades, and may contribute to behavioral abnormalities in prodromal AD. Indeed, aggregation of hyperphosphorylated tau in the LC coincides with the emergence of non-cognitive behavioral impairments commonly observed in prodromal AD that are consistent with LC dysfunction [33, 35, 36]. Lower LC integrity as measured with MRI contrast has been linked with depression, sleep disorders, and impaired cognition at stages when hyperphosphorylated tau is expected in the LC but few other regions [37–41]. LC post-mortem integrity and tangle load [42, 43] and BOLD activation/functional connectivity [44–48] have also been implicated in cognitive reserve and/or behavioral abnormalities, highlighting a central role for the LC in their dysregulation. One study even identified the LC as the origin of functional connectivity deficits in AD patients [49]. Separately, tau abundance in cerebrospinal fluid and positron emission tomography levels have been associated with non-cognitive behavioral impairments in healthy adults and in populations at risk for developing AD [50–54]. Together, these data suggest an association between the development of LC tau neuropathology and behavioral impairments in early AD, but a causal link remains speculative due to limitations in the ability for imaging techniques to localize and quantify pathology in small nuclei such as the LC [55].

Animal models are valuable tools for establishing mechanistic relationships between pathology and behavior in AD. These models have provided crucial information regarding the consequences of LC perturbations in the context of disease that are largely congruent with human studies. For example, neurotoxic LC lesions exacerbate AD neuropathology, neurodegeneration, inflammation, lethality, and/or cognitive impairment in transgenic rodent AD models [56–58]. Even in the absence of experimental LC lesions, noradrenergic denervation, altered NE signaling, and LC cell loss have been reported in these models [59–63], but there are important limitations to consider. Amyloidosis models generally fail to recapitulate tau pathology in the LC. Even the P301S mouse, which does accumulate hyperphosphorylated tau in the LC, employs a ubiquitous promoter and a mutant form of tau that causes frontotemporal dementia that simultaneously produces tau pathology all over the brain and more closely mimics pure tauopathies [56, 64]. To isolate the effects of aberrant tau in the LC, we and others have used viral vectors to express various forms of tau specifically in the LC of mice and rats [32, 65], which triggers LC degeneration, transsynaptic tau spread, and cognitive deficits. These manipulations, however, were not performed in the context of Aβ, primarily utilized mutant forms of tau not known to occur in AD, and did not assess behaviors reminiscent of non-cognitive prodromal AD symptoms.

Our lab has previously demonstrated that the TgF344-AD rat, which overexpresses mutant human amyloid precursor protein and presenilin-1 (APP/PS1), develops endogenous, age-related accumulation of hyperphosphorylated tau in the LC prior to appreciable plaque or tangle pathology in the forebrain [63]. This model provides a unique resource for understanding the consequences of hyperphosphorylated tau in the LC in the earliest phases of AD because the tau pathology (1) is triggered by AD-causing mutant Aβ, (2) is endogenous (i.e. there is no human tau transgene), and (3) follows a spatial pattern of appearance reminiscent of human disease. Dysregulation of the noradrenergic system in TgF344-AD rats is evidenced by forebrain denervation and reduced NE levels in the absence of frank neuronal loss. Behavioral characterizations of these rats have shown impaired learning and memory, hyposmia, and anxiety-like phenotypes [63, 66–69], and we have shown that chemogenetic LC activation rescues reversal learning deficits [63]. However, few studies have comprehensively tested this strain on behaviors influenced by the LC along disease progression, beginning with the appearance of tau in the LC (~6 months) and at more advanced stages, when forebrain pathology is present (~12 months).

The goals of this study were three-fold: (1) to expand the characterization of TgF344-AD rats across a battery of AD- and LC-relevant behaviors at time points where hyperphosphorylated tau in the LC is the only detectable AD-like neuropathology (6 months) and when Aβ and tau pathology is evident in the forebrain (12 months), and (2) to assess whether viral-mediated expression of wild-type (WT) human tau in the LC exacerbates AD-like behavioral phenotypes and pathology.

## Materials and Methods

### Animals

TgF344-AD rats hemizygous for the *APPsw/PS1ΔE9* transgene and WT littermates were housed in the animal facilities at Emory University. Rats were housed in groups of 2-3 on a 12-h light/dark cycle (lights on at 7:00 am) and given *ad libitum* access to food and water except during behavioral testing or otherwise specified. All procedures were approved by the Institutional Animal Care and Use Committee of Emory University.

### Stereotaxic Injections

At two months of age, rats underwent stereotaxic, sterile-tip surgery. Rats were anesthetized with isoflurane and given meloxicam or ketoprofen (2 mg/kg or 5 mg/kg, s.c., respectively) prior to receiving bilateral injections (1.3 ul/hemisphere) of either AAV9-PRSx8-hTau(WT)-WPRE-SV40 or AAV9-PRSx8-mCherry-WPRE-rBG control virus targeting the LC (AP: −3.8, ML: +/−1.2mm, DV: −7.0mm from lambda with a 15 degree downward head-tilt). Viral expression was driven by the noradrenergic-specific PRSx8 promoter [70]. Following the infusion, the injection syringe was left in place for 5 min before being moved dorsally 1mm and waiting 2 additional min to ensure diffusion of the virus at the site of injection. Behavioral assays were conducted 4 or 10 months following surgery (i.e. at 6 or 12 months of age).

### Behavioral Assays

#### General

A variety of assays were chosen to assess changes in behaviors commonly associated with prodromal and later stage AD including arousal (sleep latency), general activity levels (23-h locomotor activity), anxiety-like behaviors (open field, elevated plus maze, and novelty-suppressed feeding), passive coping that may reflect aspects of depression (forced swim test), and learning and memory (Morris water maze, cue- and context-dependent-fear conditioning). Rats were tested on behavioral tasks in the following order, from least to most stressful/invasive: sleep latency, 23-h locomotor activity, open field, elevated plus maze, forced swim task, Morris water maze, novelty-suppressed feeding, and fear conditioning. Rats were singly-housed 3-7 days before sleep latency and subsequently re-paired following completion of the 23-h locomotor test. Behavioral tests were separated by at least 1 day. All behavioral tests were conducted during the light phase under white light, unless otherwise reported.

#### Sleep latency

Approximately 2-3 hrs following lights on, rats were gently handled for 2-3 s and placed back into their home cage for 4 h and recorded with video cameras mounted above the home cage. Latency to fall asleep was quantified as the duration of time (min) until the rat had its first sleep bout. Sleep bouts were defined as periods of time that rats exhibited a sleep position for at least 75% of a 10 min time period that began with at least 2 mins of uninterrupted sleep, as previously described [71], which our lab has validated with EEG [72].

#### 23-h locomotor activity

Rats were placed into a clean cage with free access to food and water for 23 h. Cages were surrounded with 11.5 × 20” Photobeam Activity System-Home Cage infrared beams to measure locomotor activity. Total number of ambulations were binned in 30-min intervals over the 23 h testing period. Testing for all cohorts began between 9:15-10:15 am. Analyses were performed on novelty-induced locomotion, defined as occurring within the first 30 min interval, locomotion during the light and dark phases, and total locomotion over the entire 23-h period.

#### Open field

Rats were individually placed into the center of a 39” outer diameter circular arena with plastic floors and 12” gray plexiglass walls and allowed to freely explore the arena for 5 min. Activity was measured and recorded with a ceiling-mounted camera and TopScan software. Analyses focused on the duration of time spent in the inner 50% of the circle area, total velocity, and latency to exit the inner circle. Time spent in the inner circle is interpreted as lower anxiety-like behavior.

#### Elevated plus maze

Rats were individually placed into the center of an elevated plus maze (Harvard Apparatus) with two arms enclosed by 50 cm high walls and two open arms with no walls (each 50 cm × 10 cm) and allowed to freely explore for 5 min. Activity was measured and recorded with a ceiling-mounted camera and TopScan software. Rats were tested during the light phase under red lighting. Analyses focused on the time spent in the open and closed arms and total velocity. Time spent in the open arms is interpreted as lower anxiety-like behavior.

#### Forced swim test

Rats were individually placed in a clear plastic cylinder filled two-thirds with water at 25°C for 10 min. Behavior was recorded by video cameras, and immobility, defined as when the rats made only those movements necessary to keep their heads above water, was manually scored using BORIS software [73]. Immobility is indicative of passive coping behavior and was calculated using the average of scores from two blinded experimenters.

#### Morris water maze

##### Acquisition

Rats underwent four training sessions in which they were placed in a different quadrant of a 75” diameter plastic circular arena surrounded by several extra-maze cues and filled with opaque water, with a hidden platform just below the surface of the water. The platform remained in the same location throughout the acquisition sessions. Rats remained in the water until locating the platform or 60 s had elapsed, whichever came first. If 60 s elapsed, the rat was placed on the platform for 10 s. Rats completed four training sessions every day for four consecutive days. Behavior was tracked and recorded by a ceiling-mounted camera and TopScan software. Analyses were performed on distance travelled, latency to find the hidden platform, and total velocity averaged across training sessions for each day.

##### Acquisition Probe

The day following the final acquisition training day, rats were placed into the circular arena with no platform for 60 s. Analyses were performed on time spent and distance travelled in the quadrant of the maze where the platform was located during acquisition training.

##### Reversal Training

Rats underwent the same training they received during the acquisition training, but the hidden platform was moved to a different quadrant.

##### Reversal Probe

Following reversal training, rats underwent a reversal probe under identical conditions to the acquisition probe.

#### Novelty-suppressed feeding

Rats were food-deprived for 24 h and then placed in a 24 × 12 × 12” rectangular arena that contained a pre-weighed food pellet at the center of the arena during the light phase under red lighting. The session ended when the rat picked up and bit into the pellet, or when 15 min elapsed, whichever came first. Rats were subsequently placed into their home cages with the same pellet. The primary outcome measure was latency to eat in the novel environment. Shorter latency to eat the pellet in a novel environment was interpreted as lower anxiety-like behavior. To control for the possibility that there were group differences in overall hunger levels or motivation to consume food, latency to eat in the home cage immediately following the experimental session, and the amount of pellet consumed after 1 h was recorded.

#### Fear conditioning

Testing was performed over three consecutive days. After each session, rats were returned to their home cage. Freezing levels for each session were recorded with FreezeFrame software.

##### Training

For the training phase, rats were placed in a sound-enclosed metal chamber illuminated with a single light with stainless-steel bar flooring capable of delivering footshocks. Rats were allowed to habituate to the chamber for 3 min which was followed by a 2000 Hz, 80 dB tone for 20 s that co-terminated with a 2 s, 0.5 mA shock. This procedure was repeated three times with inter-tone intervals set at 60 s. Chambers were cleaned with RB10 between sessions.

#### Contextual fear conditioning

To test memory for the shock-context pairing, 24 h after Training rats were placed in the original chamber where they received training for 8 min in the absence of tone or footshock.

#### Cued fear conditioning

To test memory for the association between the cue-shock pairing, 24 h after contextual fear conditioning rats were placed in a novel chamber with a different texture (metal grid flooring) and odor (chambers cleaned with 70% ethanol before and after each session). After 1 min, rats were played the same tone from the training sessions a total of three times, with 60 s intertrial intervals in the absence of footshocks.

Primary outcome measures were freezing levels throughout contextual and cued fear conditioning. Higher freezing levels are indicative of better memory for the association between the original context and/or cue-shock pairing.

### Tissue Preparation & Immunohistochemistry

Following the conclusion of behavioral testing, rats were overdosed with isoflurane and perfused with ice cold potassium phosphate-buffered saline (KPBS) followed by 4% paraformaldehyde. Brains were removed and stored overnight in 4% PFA before being transferred to 30% sucrose until sectioning. Brains were sectioned at 40 um and stored in cryoprotectant until staining.

All antibody information can be found in Supplemental Table 1. Sections from the LC were stained with anti-norepinephrine transporter (NET) and anti-AT8 or dsRED antibodies to confirm viral expression. Sections from the hippocampus were stained for AT8 and 4G8 immunoreactivity to assess AD-related pathology. Both LC and hippocampal sections were stained with anti-ionixed calcium binding adaptor molecule 1 (IBA1) and anti-glial fibrillary acidic protein (GFAP) antibodies to mark microglia and astrocytes, respectively. Hippocampal sections were also stained with anti-NET antibody to assess LC innervation. Briefly, sections were washed 3×5 min in 0.1 M PBS and then incubated for 1 h at room temperature in blocking solution (2% normal goat serum, 1% bovine serum albumin in 0.1% Triton PBS). Sections were then incubated for 24 h at 4 degrees Celsius in blocking solution with a 1:1000 dilution of primary antibody. Sections were subsequently washed 3×5 min in PBS and incubated for 1 h at room temperature in blocking solution with an appropriate secondary antibody (1:500 dilution of goat anti-rabbit 568 for IBA1 and GFAP in the hippocampus, and dsRED, goat anti-rabbit 633 for IBA1 and GFAP in the LC, goat anti-mouse 488 for AT8, 4G8, and NET). Sections were washed 3×5 min in PBS, mounted on SuperFrost+ slides, and coverslipped with DAPI+fluoromount

### Image Analysis

For each rat, 2-3 brain sections were stained, analyzed, averaged for each animal, and subsequently averaged with animals of the same age, genotype, and virus to obtain group means. All image analysis was performed in ImageJ by blinded experimenters. In hippocampal regions a threshold was applied (Otsu), and a region of interest (ROI) of a standard size across sections (95381 pixels^2^) was used to count AT8-, IBA1-, and GFAP-positive cells in CA1, CA3, and the dentate gyrus (DG) using the analyze particles function. Plaques were analyzed as a percent area in a hippocampal subregion following thresholding. NET images were processed and analyzed as previously described [63, 74]. Briefly, the FeatureJ Hessian plugin was used to process the image, selecting the largest eigenvalue of hessian tensor, absolute eigenvalue comparison, and a smoothing scale of 0.5. An ROI of standard size (2970 pixels^2^) was used to assess the mean and standard deviation background fluorescent intensity. Two lines that were perpendicular in orientation and of standard length (208.75 pixels) were used to find peaks using the plot profile and find peaks function, which corresponded to NET fibers. Peaks were defined as fluorescent intensity values greater than the mean plus two standard deviations above background. For the LC, rolling-ball background subtraction was used to mitigate autofluorescence before quantifying IBA1 and GFAP-positive cells as described for the hippocampus. Since the size of the LC varied per section, IBA1 and GFAP-positive cells were normalized and expressed as the number of positive cells for either marker per area.

### Statistics and Analysis

All data are reported as mean ± SEM and were analyzed using a 3-way ANOVA with age, genotype, and virus as factors. When main effects were present, post-hoc analyses with Šidák correction were performed between groups differing by a single factor (age, genotype, or virus) with statistical significance set at α = 0.05. Significant statistics are presented in-text, while all statistics for behavior and immunohistochemistry are presented in Supplemental Tables 2 and 3, respectively. Statistical analysis was performed and data were graphed using Gaphpad Prism. For 23-h locomotion, Morris water maze, and fear conditioning, initial analyses considered interval/time period/training day as a factor. The area under the curve function in GraphPad prism was used to compare across ages with a 3-way ANOVA for novelty-induced, light, and dark phase in the 23-h locomotion paradigm, Morris water maze acquisition and reversal acquisition, and cued and contextual fear condition training and test days.

## Results

### Confirmation of viral expression

We confirmed robust expression of WT hTau and the mCherry control virus in the LC of WT and TgF344-AD rats at 6 and 12 months of age (Fig. 1). There was a main effect of genotype of AT8 fluorescence levels (F[1,34] = 4.857, p = 0.0344), where WT hTau injected TgF344-AD rats had elevated AT8 staining at both ages (Supplemental Fig. 1). As previously reported [63], the LC of mCherry-infected TgF344-AD rats did not stain positive utilizing the AT8 antibody, but robust AT8 immunoreactivity was observed in both WT and TgF344-AD rats that received the WT hTau virus (Fig. 1).

**Fig. 1.**
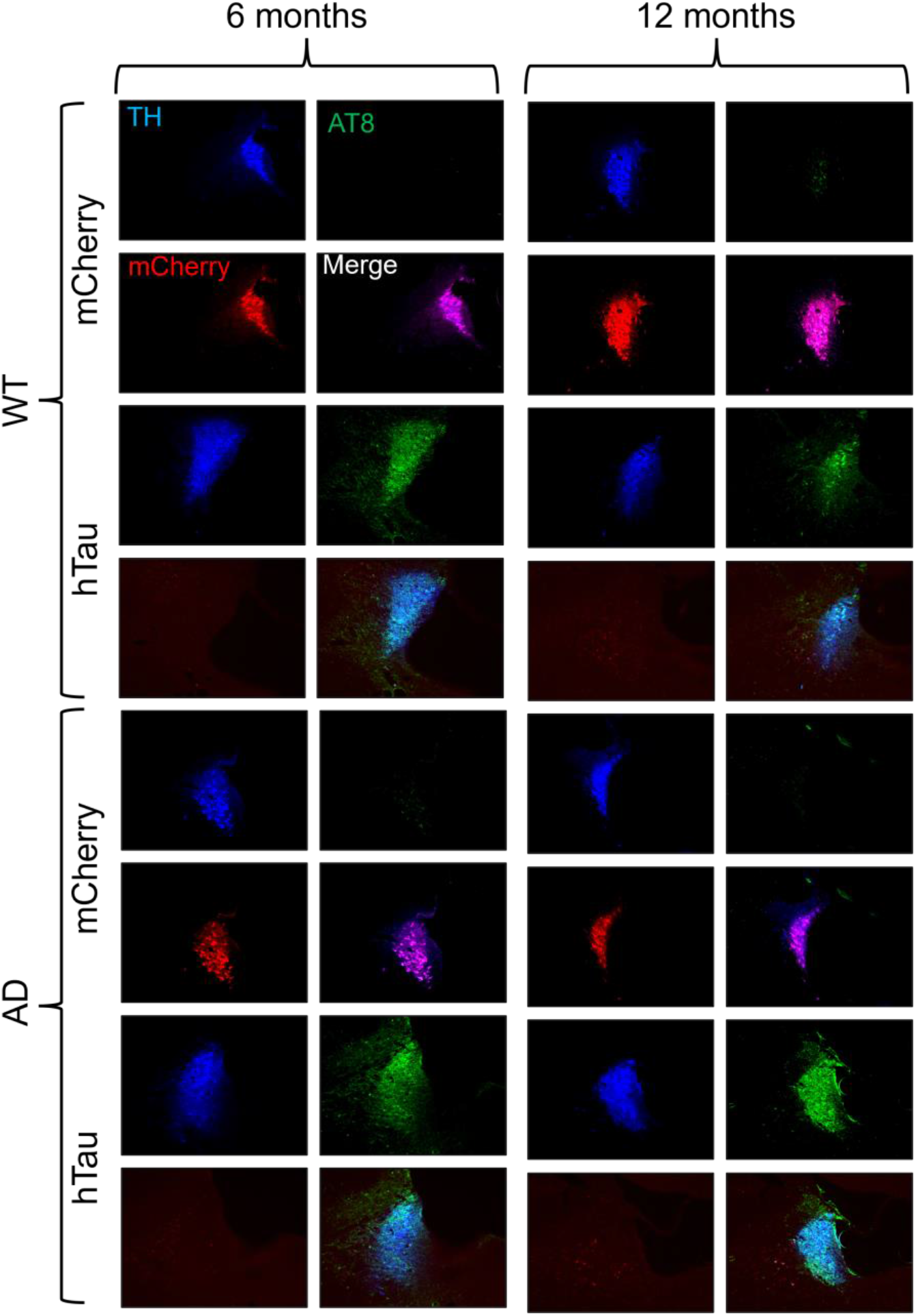
Verification of viral-mediated hTau and mCherry expression in the rat LC. Wild-type (WT) and TgF344-AD (AD) rats were injected bilaterally into the LC with AAV-PRSx8-hTau (hTau) or AAV-PRSx8-mCherry (mCherry) at 2 months of age, and assessed for hTau and mCherry expression by immunohistochemistry at 6 months or 12 months. Shown are representative 20x immunofluorescent images for the LC marker tyrosine hydroxylase (TH; blue), the hyperphosphorylated tau marker AT8 (green), mCherry (red), and merged signals.

### General Arousal and Locomotion

#### Sleep Latency

Sleep disturbances are common in AD, and often emerge coincident with tau pathology in the LC [24–26, 35]. We first sought to determine whether age, genotype, virus, or their interactions influenced wakefulness/arousal by measuring latency to fall asleep after gentle handling, and found a main effect of age (F[1,72] = 9.284, p = 0.0032), where older animals took longer to fall asleep (Fig. 2A). There was also an age × genotype × virus interaction (F[1,72] = 4.218, p = 0.0436) (Fig. 2A); at 12 months, hTau virus appeared to increase sleep latency in WT rats and decrease sleep latency in TgF344-AD rats compared to their mCherry-expressing counterparts, but no post hoc tests were significant.

**Fig. 2.**
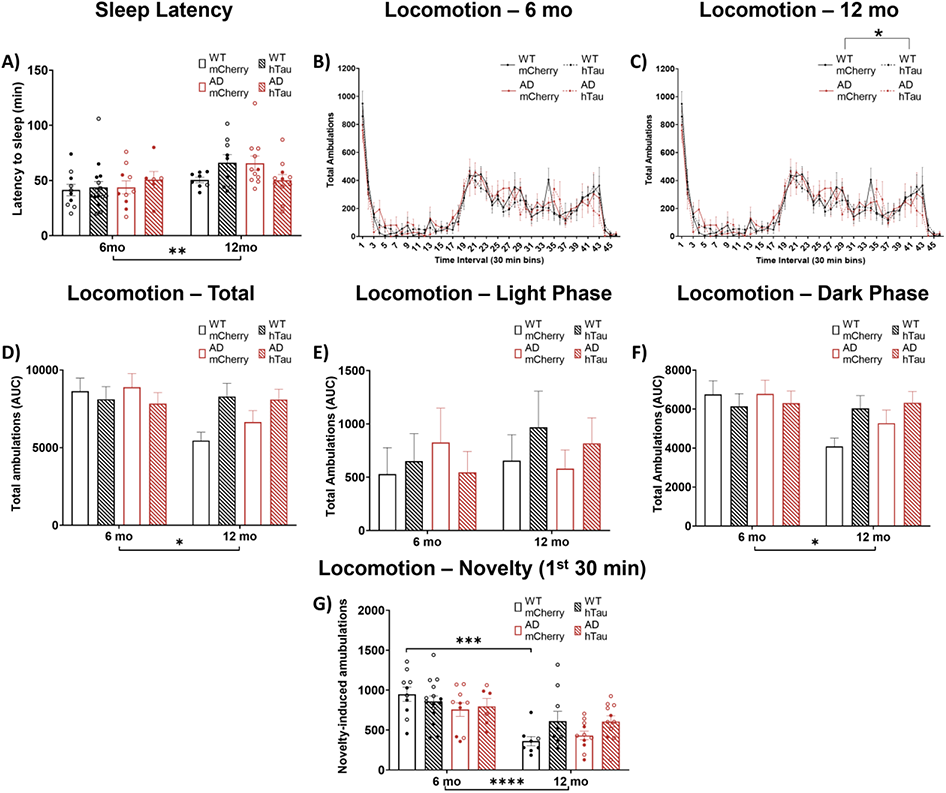
Assessment of arousal and locomotor activity. Wild-type (WT, black)) and TgF344-AD (AD, red) rats were injected bilaterally into the LC with AAV-PRSx8-hTau (hTau, hashed) or AAV-PRSx8-mCherry (mCherry, solid) at 2 months of age, and assessed for sleep latency and locomotor activity at 6 months or 12 months. Shown are latency to fall asleep after gentle handling (A), time course of 23-h locomotor activity in 6 (B) and 12 (C) month rats, total ambulations (D), light phase ambulation (E), dark phase ambulation (F), and novelty-induced locomotion during the first 30 min of the test (G). *p<0.05, **p<0.01, ***p<0.001, ****p<0.0001. Males are represented by closed circles, females by open circles.

#### 23-h locomotion

We next used chambers equipped with photobeams to assess overall arousal and locomotor activity patterns over a full day. There was a main effect of time on locomotor activity in both 6-(F[11.84,437.9] = 31.97, p < 0.0001) and 12-month rats (F[10.64,372.3] = 21.61, p < 0.0001), indicating that rats had normal circadian rhythms across the light/dark cycle. While there were no main effects of genotype, virus, or an interaction on ambulations in 6-month animals (Fig. 2B), we did detect a main effect of virus (F[1,35] = 5.88, p = 0.0206) and a virus × time interaction (F[45,1575] = 1.427, p = 0.0339) on locomotor activity in 12-month animals (Fig. 2C). We next performed an area under the curve analysis to include age as a variable. A main effect of age (F[1,72] = 4.592, p = 0.0355) and an age × virus interaction (F[1,72] = 6.304, p = 0.0143) was apparent for locomotion over the full 23-h period (Fig. 2D). Parsing the data by light/dark phase, there were no main effects during the light cycle (Fig. 2 E), but there was a main effect of age (F[1,72] = 4.821, p = 0.0313) and an age × virus interaction (F[1,72] = 4.435, p = 0.0387) during the dark period (Fig. 2F). While no comparisons survived post-hoc correction, there was a consistent pattern for 12 month mCherry rats of both genotypes to show decreased locomotor activity compared with 6 months (Fig. 2D), whereas 12-month hTau rats either maintained or even increased ambulations compared to the younger rats. Finally, we assessed novelty-induced locomotion (Fig. 2G), defined as the number of amublations within the first 30 min following placement in the chambers, and found a main effect of age (F[1,72] = 31.87, p < 0.0001) and an age × virus interaction (F[1,72] = 3.981, p = 0.0498). Older rats generally moved less than younger animals, but only WT mCherry injected rats met statistical significance (t[72] = 4.784, p = 0.0001).

### Anxiety- and active/passive coping-like behavior

#### Open field

We investigated anxiety-like behavior, which can be induced by augmenting LC activity [75–79], using a battery of canonical tests beginning with the open field test. There was a significant main effect of genotype on percent of time spent in the inner 50% of an open field (F[1,73] = 4.651, p = 0.0343), which was mainly driven by younger TgF344-AD rats that spent less time in the inner area (Fig. 3A), indicative of increased anxiety-like behavior [80]. There was also a significant effect of age on total distance traveled in the open field (F[1,73] = 14.57, p =0.0003), where older animals moved less compared with their younger counterparts (Fig. 3B), similar to some results we obtained in the locomotor activity experiment (Fig. 1).

**Fig. 3.**
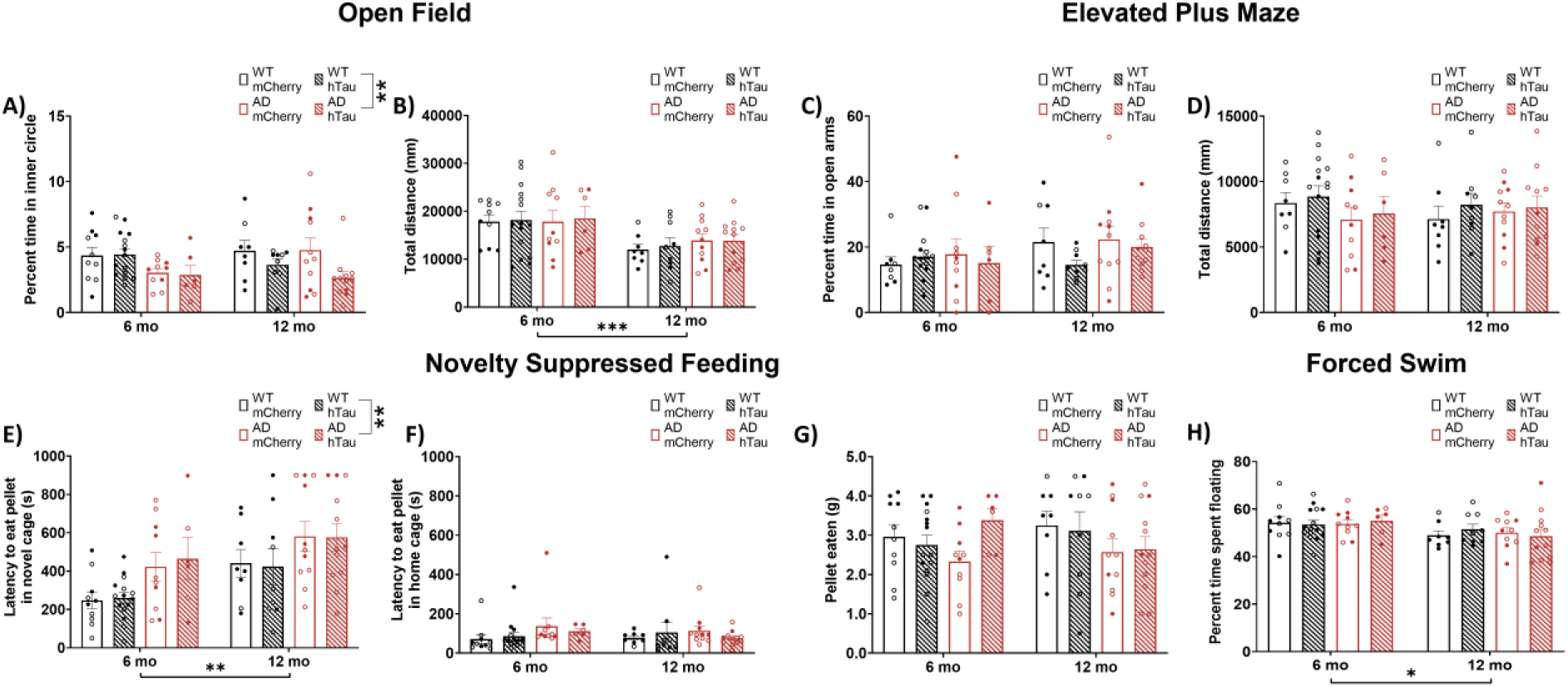
Assessment of anxiety-like behaviors and coping strategies. Wild-type (WT, black) and TgF344-AD (AD, red) rats were injected bilaterally into the LC with AAV-PRSx8-hTau (hTau, hashed bars) or AAV-PRSx8-mCherry (mCherry, open bars) at 2 months of age, and assessed for open field, elevated plus maze, forced swim test, and novelty-suppressed feeding performance at 6 months or 12 months. Shown is percent time in the inner portion (A) and total distance travelled (B) in the open field test, percent time in the open arms (C) and total distance travelled (D) in the elevated plus maze, latency to eat in a novel (E) or home/familiar (F) cage and amount of food consumed (G) in the novelty-suppressed feeding test, and time spent floating (H) in the forced swim test. *p<0.05, **p<0.01, ***p<0.001. Males are represented by closed circles, females by open circles.

#### Elevated plus maze

We next tested rats on the elevated plus maze. There were no main effects on percent time spent in the open arms (Fig. 3C) or distance traveled (Fig. 3D).

#### Novelty-suppressed feeding

Novelty-suppressed feeding (NSF) pits a hungry animal’s appetitive drive against its fear of exposure in unfamiliar open spaces, and longer latencies to eat are interpreted as increased anxiety-like behavior [81, 82]. Importantly, our lab has recently shown this task to be especially sensitive to dysregulation of LC-NE transmission [82]. There were significant main effects of age (F[1,73] = 9.9691, p = 0.0026) and genotype (F[1,73] = 11.12, p = 0.0013), where it took longer for older and TgF344-AD animals to eat the food pellet in a novel cage (Fig. 3E). To control for possible differences in hunger, we recorded the latency to eat a food pellet in the home cage (Fig. 3F) and amount of pellet eaten within 1 h (Fig. 3G). There were no main effects of age, genotype, or virus on either measure. These data are consistent with increased anxiety-like behavior in TgF344-AD rats and in older animals in general.

#### Forced swim test

The forced swim test is thought to reflect passive (floating) vs active (struggling or swimming) behavior under conditions of inescapable stress [83, 84]. There were no main effects on percent of time struggling or swimming in the forced swim test. There was a significant effect of age (F[1,72] = 6.904, p = 0.0105) on percent of time spent floating whereby older animals floated less than younger animals (Fig. 3H).

### Learning and Memory

#### Morris water maze

We have previously reported that 6- and 16-monthTgF344-AD rats exhibit modest deficits in acquisition of spatial learning in the Morris water maze, but have profound impairments in reversal learning [63]. We again employed this task to assess hippocampal-dependent learning and memory in WT and TgF344-AD rats with and without hTau expression in the LC. During acquisition, a main effect of day on distance travelled to find the hidden platform was apparent in 6-(F[2.759,102.1] = 44.77, p<0.0001, Fig. 4A) and 12-month animals (F[2.173,78.23] = 70.02, p<0.0001, Fig. 4B), indicating that animals of both ages were able to learn the location of the hidden platform with training. 6-month animals also displayed a significant effect of genotype (F[1,37] = 6.08, p = 0.0184), with impaired learning in TgF344-AD rats (Fig. 4A). For 12-month animals, a main effect of virus (F[1,36] = 5.618, p =0.0233) indicated that hTau expression was associated with reduced performance (Fig. 4B). Collapsing across training days and factoring in age revealed no main effects of age, genotype, virus, or interactions on distance travelled (Fig. 4C). For the probe trial, there was a main effect of age (F[1,73] = 5.336, p = 0.0237), where older animals spent less time in the target quadrant compared with younger animals (Fig. 4D).

**Fig. 4.**
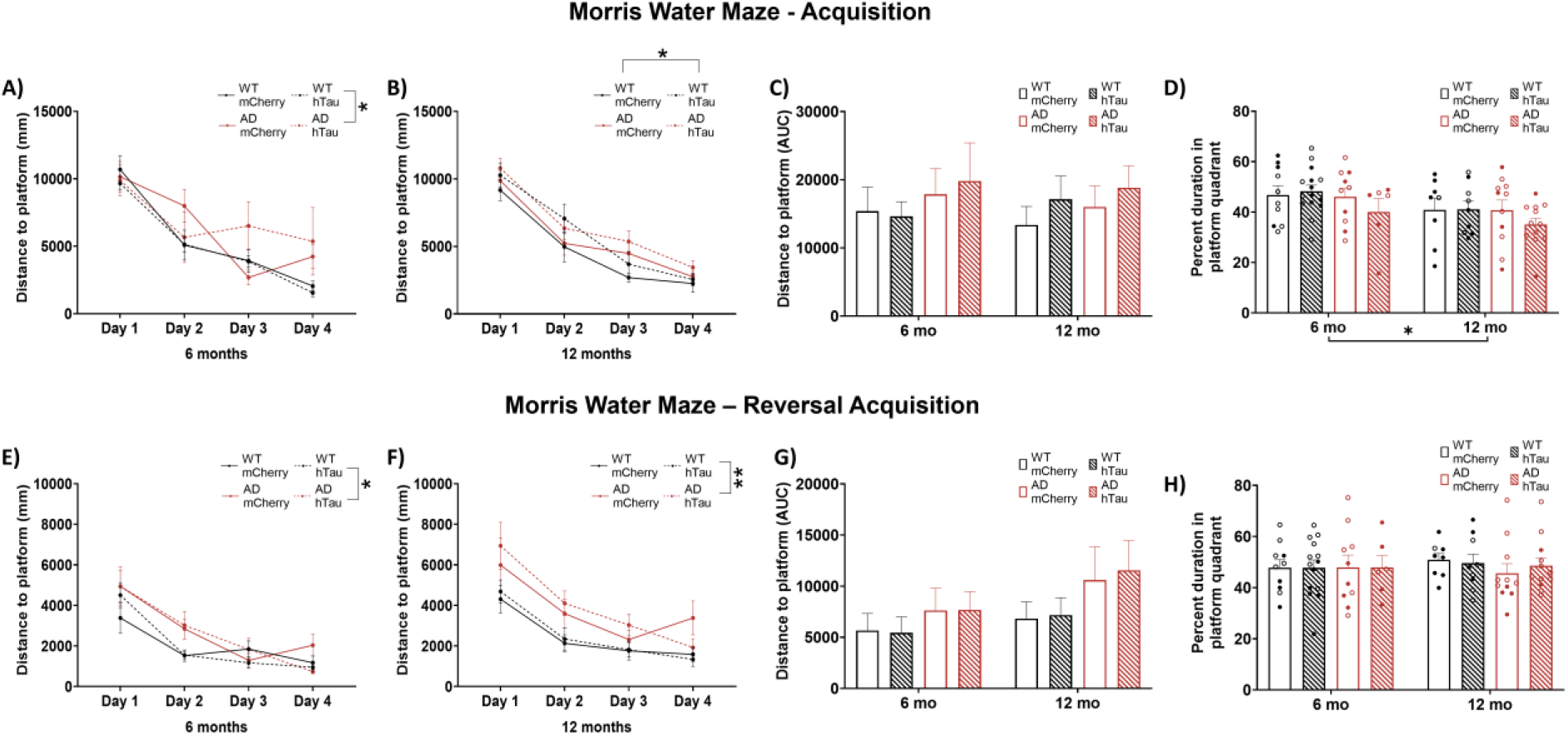
Assessment of hippocampal-dependent spatial learning and memory in the Morris water maze. Wild-type (WT, black) and TgF344-AD (AD, red) rats were injected bilaterally into the LC with AAV-PRSx8-hTau (hTau, hashed) or AAV-PRSx8-mCherry (mCherry, solid) at 2 months of age, and assessed for performance in the Morris water maze at 6 months or 12 months. Shown are total distance to find the hidden platform by day at 6 months (A) and 12 months (B), total area under the curve distance collapsed across days at both ages (C), percent time spent in target quadrant during the probe trial (D), total distance during reversal learning at 6 months (E) and 12 months (F), total area under the curve distance during reversal learning collapsed across days at both ages (G), and percent time spent in target quadrant during the probe trial after reversal learning (H). *p<0.05, **p<0.01. Males are represented by closed circles, females by open circles.

For reversal acquisition, main effects of day were again present in both 6- and 12-month animals on distance travelled to find the hidden platform (6 month: F[1.793,66.33] = 35.49, p<0.0001; 12 month: F[3,108] = 22.96, p<0.0001, Fig. 4E & F), with rats improving over training. Consistent with our previous study, reversal learning was impaired in TgF344-AD rats (main effect of genotype at 6 months, F[1,37] = 4.999, p = 0.0315, Fig. 4E) and 12-months, F[1,36] = 9.539, p = 0.0039, Fig. 4F). After collapsing days, a trend towards a main effect of genotype (F[1,73] = 3.363, p = 0.0708, Fig. 4G) was noted without any influence of age. No differences were observed in the probe trial (Fig. 4H).

#### Fear conditioning

To determine whether the impairment in spatial cognition we observed in the TgF344-AD rats in the Morris water maze extended to associative learning and memory, we performed cued and contextual fear conditioning experiments. During shock-tone pairing, a main effect of interval was apparent for both 6-(F[3.948,146.1] = 43.74, p<0.0001) and 12-month (F[3.702,133.3] = 20.58, p<0.0001) rats (Fig. 5A & B), demonstrating that all animals learned to associate the application of a tone with shock. An effect of genotype (F[1,36] = 7.184, p = 0.011) and a genotype × interval interaction (F[6,216] = 2.203, p = 0.0438) were also apparent in 12-month animals, with TgF344-AD animals freezing less during the tone/shock pairings compared with WT animals. No post-hoc tests were significant, and no differences were apparent when groups were compared across age (Fig. 5C). During subsequent exposure to the shock-paired context in the absence of the tone, there was a main effect of interval in both 6-(F[3.852,142.5] = 10.78, p<0.0001) and 12-month (F[3.265,117.6] = 9.797, p<0.0001) rats. Likewise, there was a main effect of interval present in 6-(F3.489,129.1] = 100.7, p<0.0001) and 12-month (F[3.348,120.5], = 49.29, p<0.0001) rats when presented with the shock-paired tone cue in a novel environment. These results demonstrate maintenance of the association between context/cue and shock regardless of genotype or virus group, and there were no other main effects on freezing behavior to either the context (Fig. D & E) or cue (Fig. 5G & H). Area under the curve analysis indicated a significant effect of age on freezing, where older animals froze less than younger ones under both context (F[1,73] = 6.434, p = 0.0133, Fig. 5F) and cue(F[1,73] = 4.431, p = 0.0387, Fig. 5I) conditions.

**Fig. 5.**
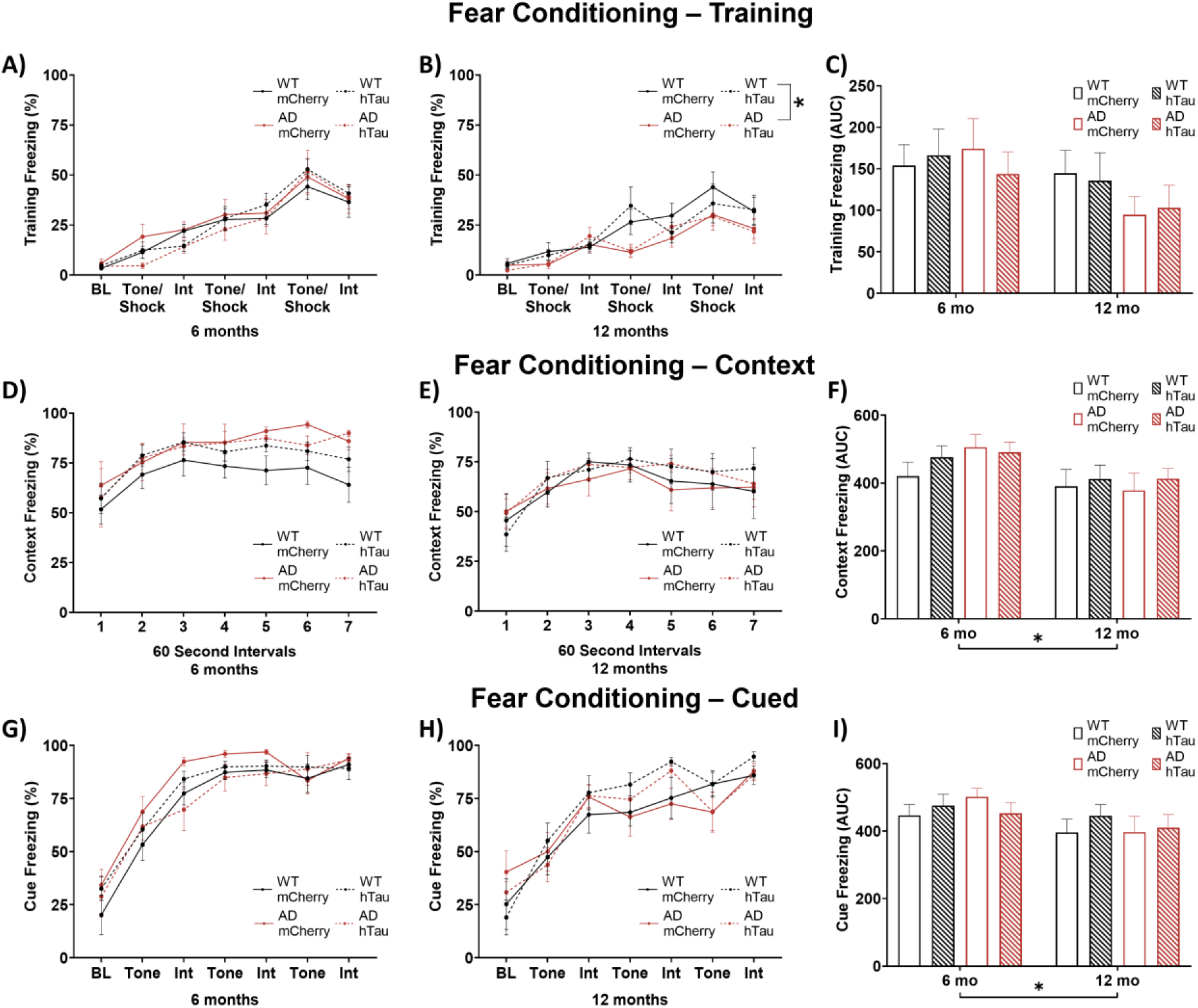
Assessment of contextual and cued fear conditioning. Wild-type (WT, black) and TgF344-AD (AD, red) rats were injected bilaterally into the LC with AAV-PRSx8-hTau (hTau, hashed) or AAV-PRSx8-mCherry (mCherry, solid) at 2 months of age, and assessed for contextual and cued fear conditioning at 6 months or 12 months. Shown is % freezing during shock-tone pairing in 6-month (A) and 12-month (B) rats, area under the curve collapsed across both ages (C), during subsequent context exposure in 6-month (D) and 12-month (E) rats, area under the curve collapsed across both ages (F), during subsequent cue exposure in 6-month (G) and 12-month (H) rats, and area under the curve collapsed across both ages (I). *p<0.05.

#### Hippocampus Pathology

AD-like neuropathology has previously been reported in the hippocampus of TgF344-AD rats [63, 66]. To expand on these analyses and determine the effect of hTau expression in the LC, we performed immunohistochemistry for Aβ (4G8), hyperphosphorylated tau (AT8), and neuroinflammation (GFAP and IBA1). Representative images of amyloid pathology in the DG, CA3, and CA1 are shown in Fig. 6A and in Supplemental Fig. 2A & B. There were main effects of age, genotype, and an age × genotype interaction on the percentage of plaque-positive area in all three hippocampal subregions (DG: Age: F[1,38] = 34.36, p < 0.0001; Genotype: F[1,38] = 92.88, p < 0.0001; Age × Genotype: F[1,38] = 34.32, p < 0.0001, Fig. 6B; CA3: Age: F[1,38] = 69.53, p < 0.0001; Genotype: F[1,38] = 101.1, p < 0.0001; Age × Genotype: F[1,38] = 69.32, p < 0.0001, Fig. 6C; CA1: Age: F[1,38] = 46.15, p < 0.0001; Genotype: F[1,38] = 73.95, p < 0.0001; Age × Genotype: F[1,38] = 47.14, p < 0.0001, Fig. 6D). Post-hoc analyses indicated 12-month TgF344-AD rats had elevated plaque burden when compared with 6-month TgF344-AD rats (DG: mCherry animals: t[38] = 4.719, p = 0.0004, hTau animals: t[38] = 7.228, p <0.0001; CA3: mCherry animals: t[38] = 8.053, p < 0.0001, hTau animals: t[38] = 8.954, p <0.0001; CA1: mCherry animals: t[38] = 6.681, p < 0.0001, hTau animals: t[38] = 7.260, p < 0.0001) and age- and virus-matched WT littermates across all hippocampal subregions (DG: mCherry animals: t[38] = 7.371, p < 0.0001, hTau animals: t[38] = 8.441, p <0.0001; CA3: mCherry animals: t[38] = 9.054, p < 0.0001, hTau animals: t[38] = 9.716, p <0.0001, CA1: mCherry animals: t[38] = 8.178, p < 0.0001, hTau animals: t[38] = 7.615, p <0.0001). There were no effects of hTau expression on plaque pathology across age or genotype.

**Fig. 6.**
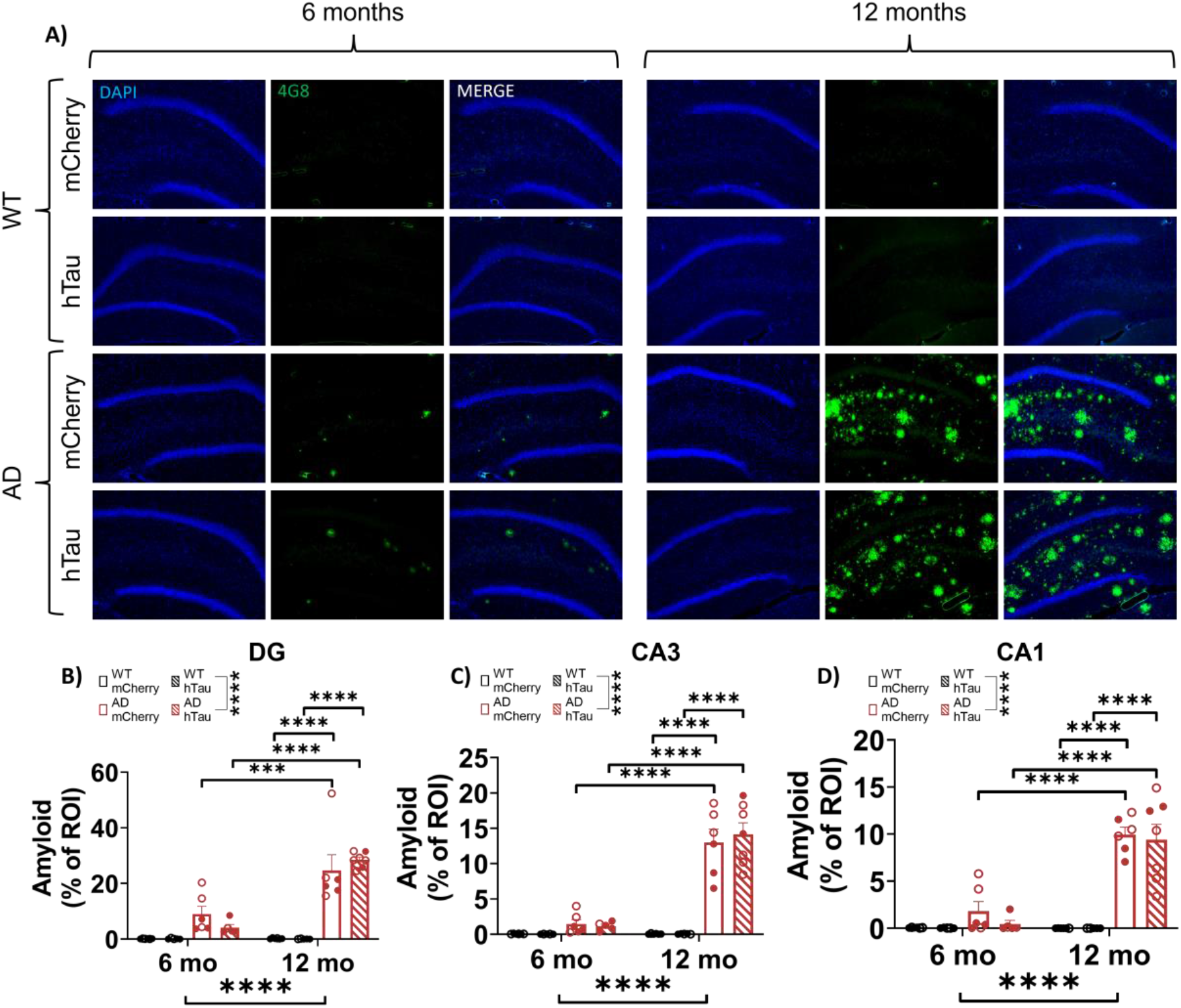
Amyloid pathology in the hippocampus. Wild-type (WT, black) and TgF344-AD (AD, red) rats were injected bilaterally into the LC with AAV-PRSx8-hTau (hTau, hashed) or AAV-PRSx8-mCherry (mCherry, solid) at 2 months of age, and assessed for 4G8 (i.e., Aβ) immunoreactivity at 6 months or 12 months. Shown are representative images of amyloid pathology in the DG subregion of the hippocampus (A), and quantification of amyloid pathology in the DG (B), CA3 (C), and CA1 expressed as % region of interest (ROI) (D). Images taken at 20x magnification. ***p<0.001, ****p<0.0001. Males are represented by closed circles, females by open circles.

To assess potential contribution of hTau expression in the LC to hippocampal tau pathology, AT8 was used to visualize hyperphosphorylated tau. Representative images of AT8 pathology in the DG CA3, and CA1 are shown in Fig. 7A and in Supplemental Fig. 3A & B. There was a main effect of genotype (F[1,33] = 4.467, p = 0.0422) in the DG, where TgF344-AD rats had higher levels of hyperphosphorylated tau pathology compared with their WT littermates (Fig. 6B), as well as an age × genotype interaction (F [1,38] = 6.305, p = 0.0164) in CA3, where older TgF344-AD rats had elevated AT8 staining compared to their WT littermates (Fig. 6C). However, for CA3, no significant differences were evident during post-hoc comparisons. In CA1, there was a main effect of age (F [1,38] = 8.648, p = 0.0055) and an age × genotype (F[1,38] = 7.596, p = 0.0089) interaction (Fig. 6D). Post-hoc comparisons revealed increased AT8+ cells in 12-month mCherry injected TgF344-AD animals compared with their 6-month WT counterparts (t[38] = 3.689, p = 0.0084). Similar to plaque pathology, there were no effects of hTau expression on AT8 immunoreactivity across age or genotype.

**Figure 7.**
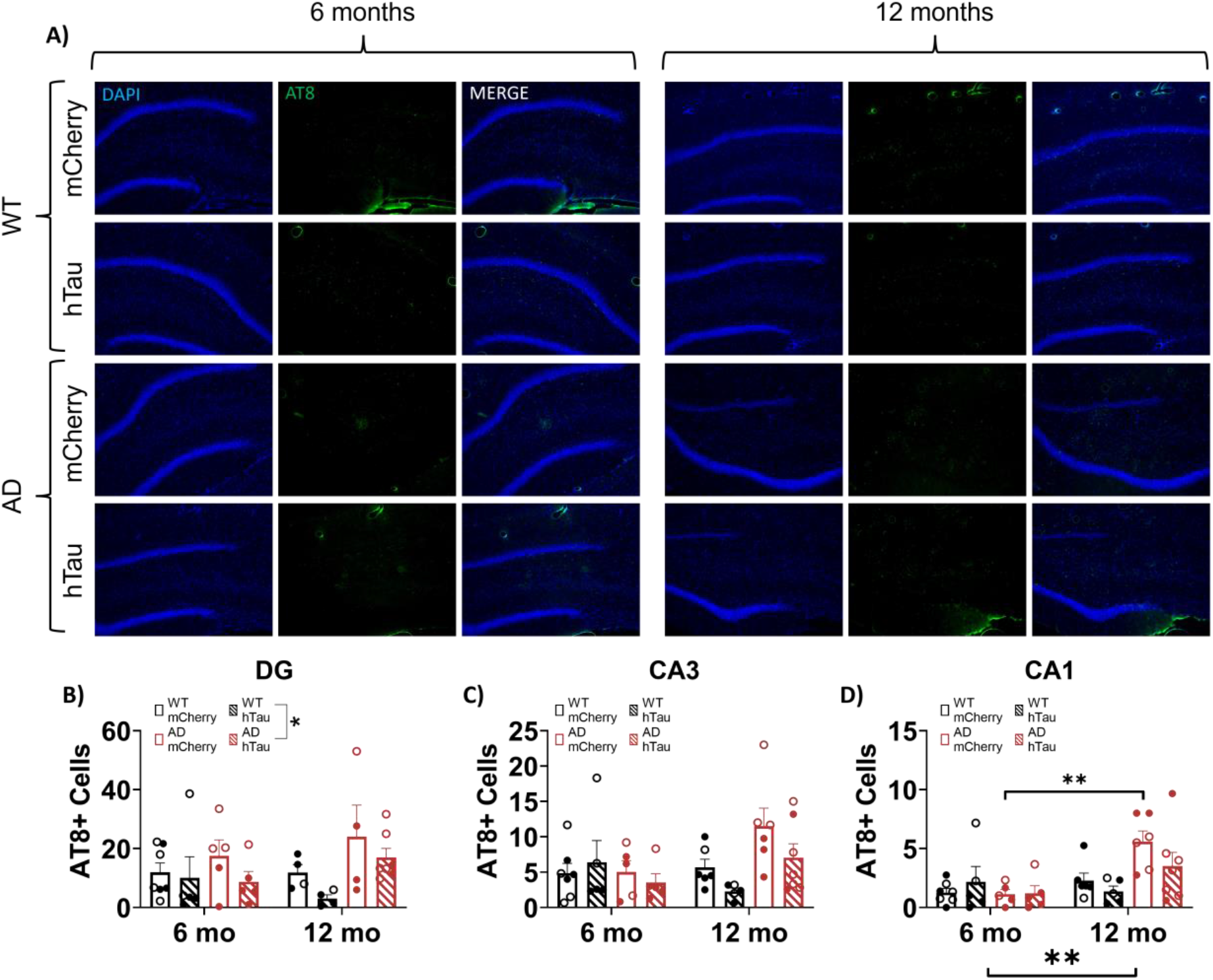
Assessment of hyperphosphorylated tau. Wild-type (WT, black) and TgF344-AD (AD, red) rats were injected bilaterally into the LC with AAV-PRSx8-hTau (hTau, hashed) or AAV-PRSx8-mCherry (mCherry, solid) at 2 months of age, and assessed for AT8 (i.e., hTau) immunoreactivity at 6 months or 12 months. Shown are representative images of hyperphosphorylated tau pathology in the DG subregion of the hippocampus (A), and quantification of hyperphosphorylated tau pathology in the DG (B), CA3 (C), and CA1 expressed as the number of AT8+ cells. (D). Images taken at 20x magnification. *p<0.05, **p<0.01. Males are represented by closed circles, females by open circles.

We next assessed neuroinflammatory astrocyte (GFAP) and microglia (IBA1) markers that are noted in AD. Representative images of GFAP staining in the DG, CA3, and CA1 are shown in Fig. 8A and in Supplemental Fig. 4A & B. There was a main effect of age, genotype, and an age x genotype interaction on number of GFAP+ cells in the DG (Age: F[1,38] = 21.56, p = <0.0001; Genotype: F[1,38] = 36.66, p <0.0001; Age x Genotype: F[1,38] = 12.04, p = 0.013, Fig. 7B), CA3 (Age: F[1,38] = 27.91, p < 0.0001; Genotype: F[1,38] = 31.43, p < 0.0001; Age x Genotype: F[1,38] = 14.11, p = 0.0006, Fig. 8C), and CA1 (Age: F[1,38] = 23.76, p < 0.0001; Genotype: F[1,38] = 5.528, p = 0.0240; Age x Genotype: F[1,38] = 4.730, p = 0.0359, Fig. 8D). Post-hoc analysis of the DG revealed elevated GFAP+ cells in 12-month mCherry (t[38] = 4.077, p = 0.0027) and hTau (t[38] = 4.204, p = 0.0018) injected TgF344-AD rats compared with their 6-month counterparts. In addition, 12-month mCherry (t[38] = 4.640, p = 0.0005) and hTau (t[38] = 5.080, p = 0.0001) injected TgF344-AD rats had elevated GFAP+ cells in the DG compared with their WT age-matched littermates. Similarly, 12-month mCherry (t[38] = 5.054, p = 0.0001) and hTau (t[38] = 4.180, p = 0.002) injected TgF344-AD rats showed increased GFAP+ cells in CA3 compared with 6-month animals. 12-month mCherry (t[38] = 5.271, p < 0.0001) and hTau (t[38] = 4.294, p = 0.0014) injected TgF344-AD rats also demonstrated elevated GFAP+ cells in the CA3 compared with WT age-matched littermates. Within CA1, 12-month mCherry injected TgF344-AD rats had more GFAP+ cells compared with 6-month TgF344-AD rats (t[38] = 5.040, p = 0.0001) and 12-month WT controls (t[38] = 3.208, p = 0.0321).

**Fig. 8.**
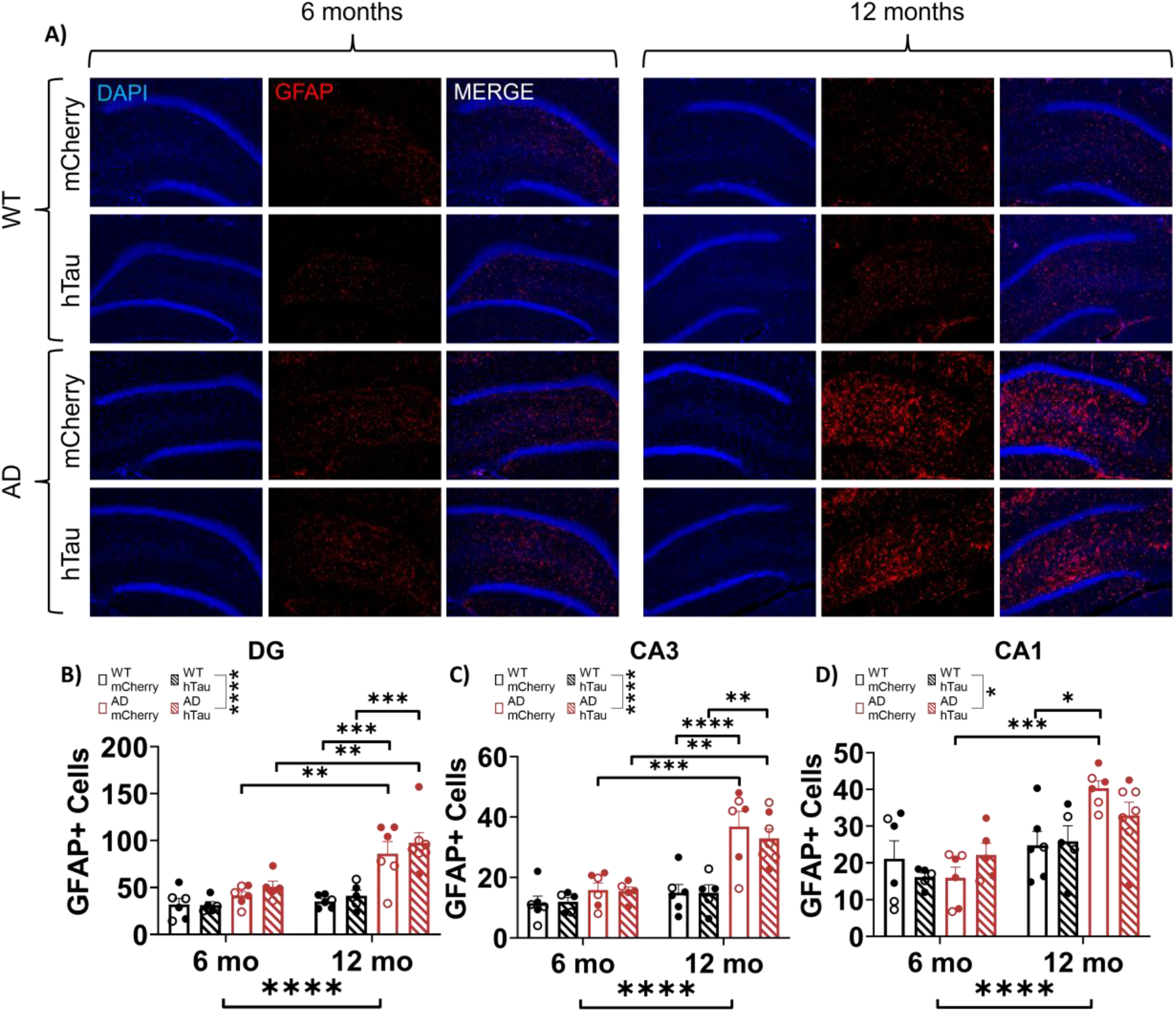
Assessment of astrocyte inflammation. Wild-type (WT, black) and TgF344-AD (AD, red) rats were injected bilaterally into the LC with AAV-PRSx8-hTau (hTau, hashed) or AAV-PRSx8-mCherry (mCherry, solid) at 2 months of age, and assessed for GFAP immunoreactivity at 6 months or 12 months. Shown are representative images of astrocyte inflammation in the DG subregion of the hippocampus (A), and quantification of astrocyte inflammation in the DG (B), CA3 (C), and CA1 expressed as the number of GFAP+ cells. (D). Images taken at 20x magnification. *p<0.05, **p<0.01, ***p<0.001, ****p<0.0001. Males are represented by closed circles, females by closed circles.

Representative images of IBA1 staining in the DG, CA3, and CA1 are shown in Fig. 9A and in Supplemental Fig. 5A and B. There was a main effect of genotype (F[1,38] = 8.596, p = 0.0057) in the DG, where transgenic animals appeared to have more IBA1+ cells than their WT littermates (Fig. 9B). There were trends towards main effects of age (F[1,39] = 3.960, p = 0.0536), genotype (F[1,39] = 3.108, p = 0.0857), and virus (F[1,39] = 3.379, p = 0.0736) on IBA1+ cells in CA3 (Fig. 9C). There was a main effect of age (F[1,37] = 24.81, p < 0.0001) on IBA1+ cells in CA1 (Fig. 9D).

**Fig. 9.**
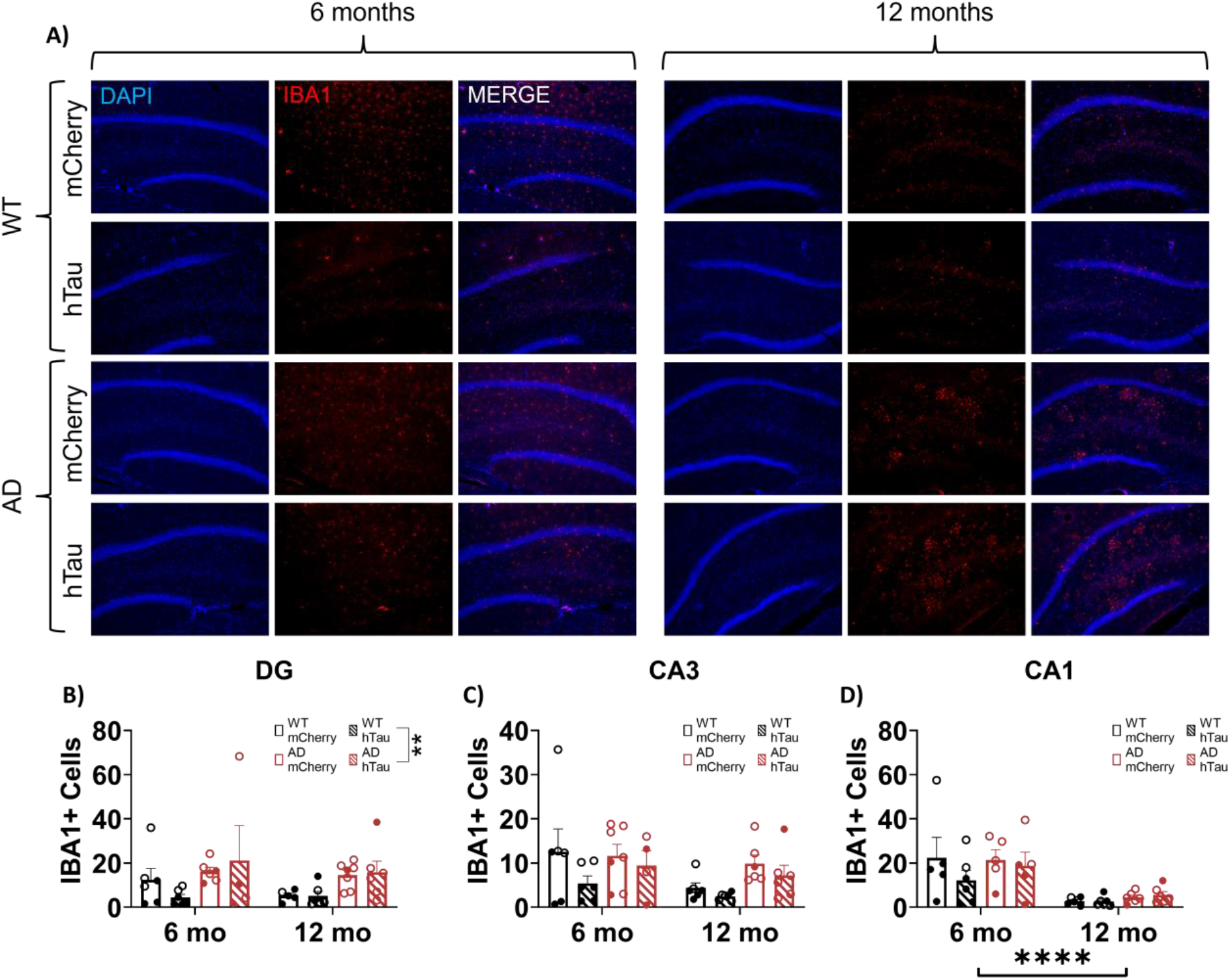
Assessment of microglial inflammation. Wild-type (WT, black) and TgF344-AD (AD, red) rats were injected bilaterally into the LC with AAV-PRSx8-hTau (hTau, hashed) or AAV-PRSx8-mCherry (mCherry, solid) at 2 months of age, and assessed for IBA1 immunoreactivity at 6 months or 12 months. Shown are representative images of microglial inflammation in the DG subregion of the hippocampus (A), and quantification of astrocyte inflammation in the DG (B), CA3 (C), and CA1 expressed as the number of IBA1+ cells. (D). Images taken at 20x magnification. Males are represented by closed circles, females by open circles.

#### Hippocampus NE Innervation

Noradrenergic projections from the LC to the hippocampus, particularly the DG, deteriorate over time in TgF344-AD rats, which contributes to cognitive impairment [63, 85]. Here, we used NET+ fiber density to assess noradrenergic innervation to the distinct hippocampal subfields. Representative images of NET staining in the DG, CA3, and CA1 are shown in Fig. 10A and in Supplemental Fig. 6A & B. There was a significant effect of genotype (F[1,38] = 16.36, p = 0.0002) in the DG (Fig. 10B). In CA3, there was a main effect of age (F [1,40] = 8.286, p = 0.0064), where older animals generally had lower NET fiber density compared to younger animals (Fig. 10C), while there were no main effects on in CA1 (Fig. 10D). These data confirm the loss of LC innervation to the hippocampus in TgF344-AD rats, with the DG exhibiting the highest susceptibility.

**Figure 10.**
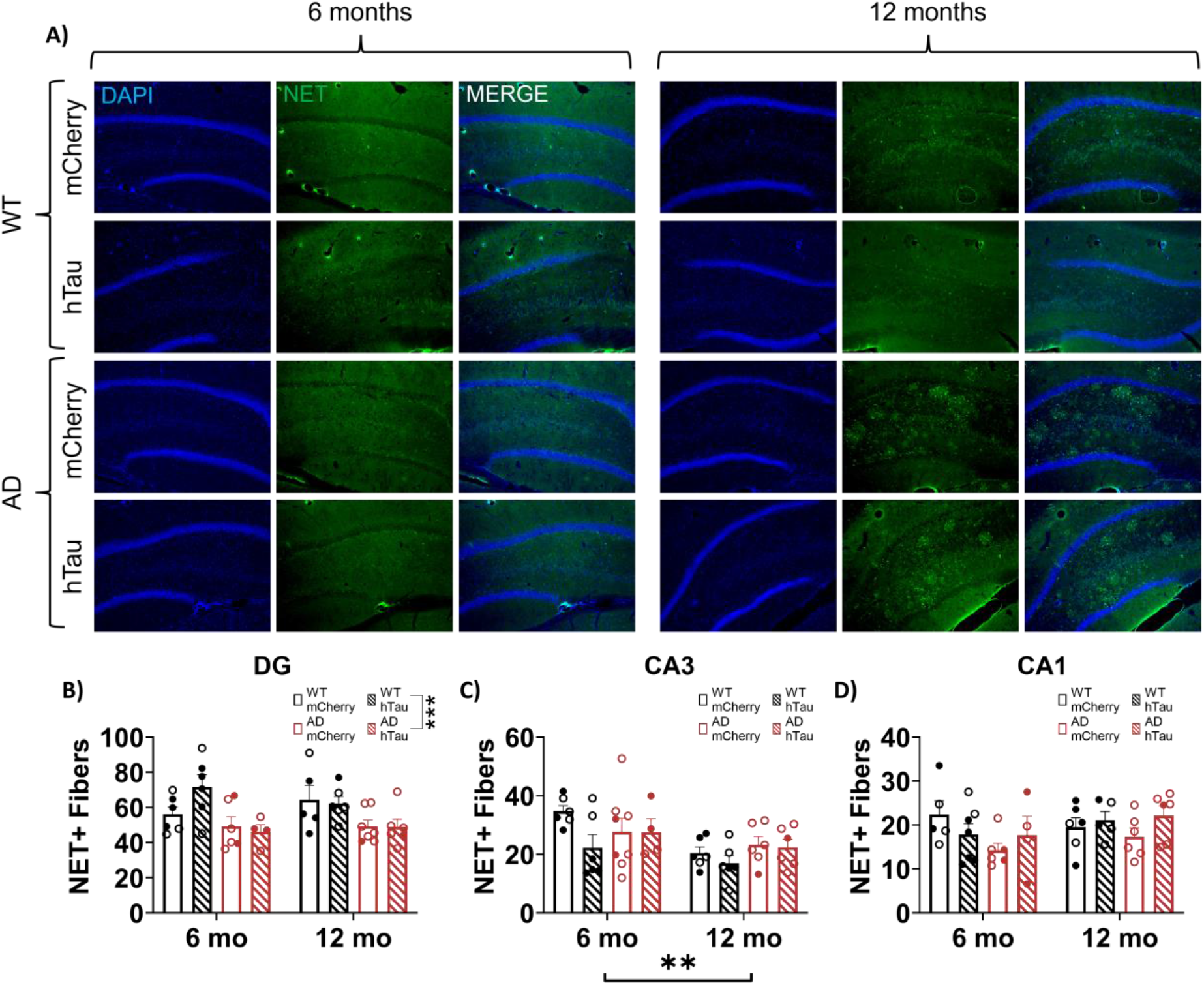
Assessment of LC fiber density in the hippocampus. Wild-type (WT, black) and TgF344-AD (AD, red) rats were injected bilaterally into the LC with AAV-PRSx8-hTau (hTau, hashed) or AAV-PRSx8-mCherry (mCherry, solid) at 2 months of age, and assessed for NET immunoreactivity at 6 months or 12 months. Shown are representative images in the DG subregion of the hippocampus (A), and quantification in the DG (B), CA3 (C), and CA1 expressed as the number of NET+ fibers. (D). Images taken at 20x magnification. **p<0.01, ***p<0.001. Males are represented by closed circles, females by open circles.

## Discussion

The LC is one of the first brain regions that accumulates hyperphosphorylated “pretangle” tau pathology at a time when notable behavioral changes begin to emerge, years or even decades prior to cognitive impairment. In this study, we set out to determine the effects of age, Aβ-triggered endogenous rat tau pathology, and virally-induced WT human tau pathology in the LC on AD-relevant behavioral phenotypes, pathology, and neuroinflammation. We observed significant effects of age and TgF344-AD genotype on behavioral and pathological markers of AD, whereas WT hTau expression in the LC was mostly inert. In general, older rats took longer to fall asleep, had reduced locomotor activity, mixed phenotypes in tests of anxiety-like behavior and passive coping, and impaired learning and memory compared with younger animals. The presence of AD-like neuropathology tended to increase anxiety-like behavior and impair cognitive performance. There are mixed reports on sex differences in behavioral phenotypes in these rats, and we were underpowered to detect differences based on sex. However, beyond differences in general locomotion, where females appeared to move more than males (see Fig. 2G and Fig. 3B, D), we did not observe obvious trends in other behavioral measures.

Changes in arousal and sleep disturbances are nearly ubiquitous in AD [24–26]. A previous study reported that 17-month TgF344-AD rats exhibited atypical EEG sleep/wake features compared to age-matched controls [86], but did not include a younger cohort for comparison. We chose to perform the sleep latency test which is an EEG-validated measure of arousal sensitive to noradrenergic manipulation [71, 72, 87]. We observed a main effect of age (older animals took longer to fall asleep) and a complex interaction between age, genotype, and virus on sleep latency of questionable physiological significance due to its modest influence. We also observed effects of age on 23-h locomotion over the total time of the task, similar to a previous report [67], and within the first 30-min. Together, results indicate that 12-month animals took longer to fall asleep but were generally less mobile. Moreover, in 12-month animals specifically, the presence of WT hTau induced an increase in locomotor activity over the entire light/dark cycle, one of the few viral-expression induced phenotypes seen here. This phenotype mimics those seen in mouse models of tauopathy [88–90], indicating these effects maybe be isoform- and/or time-specific or depend on additional influence of forebrain pathology.

Anxiety and depression are highly prevalent in prodromal populations and increase risk of developing AD [28, 29, 35, 91, 92]. When testing for anxiety-like phenotypes, we observed that 6-month TgF344-AD rats spent less time in the center of an open field compared with littermate controls, while no differences were observed in time spent in the open arms of the elevated plus maze. It has previously been reported that TgF344-AD rats show increased anxiety-like behaviors at 12, but not 2 months of age in these tasks [93]. However, these canonical tests of anxiety-like behaviors assume that rodents are motivated to explore novel environments. A lack of anxiety-like phenotypes in these tasks could be masked by differences in exploratory behavior and/or general locomotion, which we and others have observed over the course of aging and in TgF344-AD rats [67, 93]. We therefore employed the NSF task, where the main conflict is fear of a novel environment versus drive to consume food in a hungry animal. The NSF task has previously been validated as a NE-sensitive anxiety-like task by our lab using dopamine-β hydroxylase knockout mice, which lack phenotypes in canonical exploration-based anxiety paradigms [82]. We observed that both older and transgenic rats took longer to eat the food pellet in a novel environment compared with younger and age-matched WT littermates, respectively. These effects are unlikely to be mediated by differences in hunger levels because rats did not differ in the total amount of pellet eaten or latency to eat the pellet in their home cage. Therefore, accumulating evidence from our lab suggests that NSF may be particularly useful at gauging noradrenergic-specific impacts on anxiety-like behaviors.

In the forced swim task, we observed that 12-month rats spent less time spent floating than younger animals. Given the acute nature of this paradigm and the fact that older rats generally moved less in land-based locomotion assays, these results support the notion of an increase in active coping behaviors in older rats [83, 84]. A previous report demonstrated an increase in immobility in TgF344-AD rats at 12 months of age [93], which is opposite of the effects reported here. Differences may be attributed to multiple exposures to forced swim, influences of sex, or other factors known to influence coping-like phenotypes in the forced swim task [83, 94].

Spatial learning and memory deficits, which are hallmark behavioral phenotypes of AD, presented here are largely congruent with previous studies of the TgF344-AD strain, where the most robust genotype differences arise in reversal learning [63, 66]. These differences were primarily observed in acquisition of reversal learning, and were exacerbated by age in TgF344-AD rats. Interestingly, we also detected deficits in initial acquisition in 6-month TgF344-AD rats, and hTau virus induced acquisition deficits in 12-month WT rats that were reminiscent of those we observed in 6-month transgenic rats. In both of those cases (6 month Tg rats and 12 months WT rats with hTau), the main detectable AD-like neuropathology present in the brains was hyperphosphorylated tau in the LC, suggesting a causal relationship. We sought to further categorize the nature and anatomical specificity of these deficits by testing these rats on cued and contextual fear conditioning, which are associative learning tasks as opposed to spatial. Additionally, cued fear conditioning is non-hippocampal dependent, unlike both the Morris water maze and contextual fear conditioning. During training, 12-month TgF344-AD rats froze less than WT animals, which suggests a lower ability to form initial associations between the tone/shock pairing. However, genotype differences were not observed during cued or contextual fear conditioning, indicating that recall of these associations once formed was intact. Main effects of age were evident where older animals froze less to both the cue and the context than younger counterparts, suggesting a modest age-related impairment of associative memory.

In this study, we expanded on our previous work demonstrating that TgF344-AD rats have progressive noradrenergic fiber loss [63] by also showing lower NET+ fiber density in TgF344-AD rats, specifically in the dentate gyrus. Recent studies further confirm a reduction in LC innervation to the dentate gyrus [85], suggesting selective vulnerability of NE fibers within this hippocampal subfield. We also reaffirmed an age-dependent decrease in NE fibers within CA3, 4 months earlier than previously reported [63], in addition to confirming a lack of change in NE innervation to CA1. Given the loss of DBH+ fibers and reduction in NE content in the hippocampus, a loss of NET+ fibers could be indicative of compensatory mechanisms downregulating NE reuptake to increase noradrenergic signaling. Indeed, other noradrenergic compensatory changes, including enhanced β adrenergic receptor function and axonal sprouting, have been noted in TgF344-AD rats [85] and human AD/dementia cases [95, 96].

We did observe increasing levels of amyloid deposition beginning at 6 months in TgF344-AD rats across hippocampal subregions, which is slightly earlier than previously reported [66, 67]. By contrast, there were low levels of endogenous hyperphosphorylated tau across all hippocampal subregions and lack of support for transsynaptic spread of hTau from the LC. This was somewhat surprising, given the synergistic effects of amyloid and tau deposition [97]. We based our viral expression and testing paradigm based on previous literature, where pseudophosphorylated tau demonstrated transsynaptic spread and induced cognitive deficits 7 months post-injection in ~10 month old rats [32]. The viral form of tau used here (WT hTau) is a milder version from a pathological perspective compared to that reported previously (pseudophosphorylated at 14 sites), and the endogenous LC tau pathology in TgF344-AD rats is an even milder form that only reacts with the CP13 antibody (Ser202), which does not mature into more toxic species observed in the typical progression of AD [63]. Although we observed no differences in level of hTau expression within the LC between 6 and 12 month rats, it is possible that longer expression time (e.g. 14 or more months post-infusion) or initial injection of the virus into aged rats could be necessary to trigger appreciable effects on behavior and pathology. Indeed, we did observe a potential “seeding” effect, whereby TgF344-AD rats developed more AT8 pathology in the LC compared to WT littermates.

Behavioral differences between genotypes are likely attributable to development of AD-like neuropathology and inflammation over the course of aging. The effects of tau pathology on neural activity are mixed, as both hyper- and hypo-activity have been reported [98–101]. These likely depend on specific tau isoform and neuronal population, and changes in LC activity throughout the course of AD have yet to be characterized. Furthermore, the TgF344-AD rats harbor the same mutations as the APP/PS1 mouse counterpart, and APP/PS1 mice develop intraneuronal Aβ oligomers within the LC that induce noradrenergic neuron hyperactivity [102]. Given that the transgenes are expressed under the same promoter in TgF344-AD rats [66], they also likely develop Aβ oligomers. The additional accumulation of hyperphosphorylated tau within the LC could lead to a complex interaction between the two AD-like pathologies [97]. Thus, it will be important for future studies to chronical the time course of AD-like neuropathology accumulation within the LC of TgF344-AD rats and directly measure LC firing.

Changes in LC activity can also be speculated based on behavioral measures and align with the predicted accumulation and effects of neuropathology LC pathology. Generally, anxiety-like behaviors are promoted by augmented basal LC firing (3-5 Hz) [75–79]. In addition, the LC responds with phasic discharge to novelty [103, 104], and probable/confirmed AD patients show impaired novelty processing and perform worse on tasks incorporating novelty [105–107]. Similarly, aged TgF344-AD rats also demonstrate reduced ability to distinguish novel objects [66], which may be partially driven by novelty-induced anxiety-like phenotypes, as demonstrated by the NSF task. This effect is reversed in young TgF344-AD rats, which present with enhanced novelty detection that is mediated by β-adrenergic receptors [85], suggesting an augmented phasic response. Overall, results from the NSF task lend support to an age- and AD-like neuropathology-dependent alteration in LC activity in TgF344-AD rats [108]. Specifically, in young rats, enhanced signal-to-noise ratio (phasic:tonic firing) could support maintenance of novelty detection, but may lead to anxiety-like phenotypes. This effect may be reversed in 12-month rats, where signal-to-noise ratio of the LC is diminished, resulting in reduced novelty detection, and heightened basal LC activity, which also prompts expression of anxiety-like behaviors. This phenotype might be masked in other canonical tasks, such as the open field and elevated plus maze, due to differences in exploratory behavior and general locomotion, which we and others have observed over the course of aging and in TgF344-AD rats [66, 67]. We have previously reported impairments in acquisition and reversal acquisition of the Morris water maze in TgF344-AD rats [63], and here demonstrate impairments in training during fear conditioning, but only in 12-month TgF344-AD rats. However, we saw no impairments in probe trials of the Morris water maze or freezing in the contextual and cued fear conditioning paradigms, suggesting intact memory retrieval mechanisms. TgF344-AD rats present with decreased noradrenergic forebrain innervation regions early in disease [63, 85]. Spatial and associative hippocampal-dependent and independent learning and memory are supported by LC-NE release and are sensitive to noradrenergic perturbations [85, 109–111]. The loss of forebrain LC fibers could engage compensatory mechanisms, such as increased β-adrenergic receptor function, and potentially increased firing could support the maintenance of cognition in prodromal phases of AD. For example, heightened β-adrenergic receptor function facilitates extinction learning and novel object recognition [85] in younger (6-9 month) TgF344-AD rats, but these studies should be expanded to additional paradigms.

Independent of genotype effects, we observed aging effects on behavior that are suggestive of altered LC activity. 12-month rats spent less time floating in the forced swim test, indicative of an increase in active coping behaviors in response to stress. Wistar Kyoto rats, a common rodent model of depression, display increased immobility and enhanced LC activity when tested on the forced swim [112, 113]. Moreover, coping behaviors mediated by the LC may be partially dependent on galanin co-release [114], which is expressed in a high percentage of LC neurons [115]. Eliminating galanin from the LC promotes active coping behaviors in mice [114], an effect that is mimicked in older rats tested in the forced swim task. Though we cannot completely rule out frank LC cell loss over the course of aging, 16-month TgF344-AD rats do not exhibit LC degeneration compared with WT littermates [63]. Given that galanin co-expressing LC neurons are protected in AD [116], we instead theorize that changes in firing rates could reduce galanin co-release and promote active coping behaviors. Although we did not observe changes in sleep latency that would be indicative of increased LC activity, other arousal systems that have yet to be explored in these rats could compensate for LC-specific abnormalities. Alternatively, there is evidence for decreased LC activity with aging [117], but the effects may be strain-dependent. The presence of AD-like neuropathology could further modulate age-related changes in LC activity, but this has not been investigated.

Our results expand the characterization of behavior and AD-like neuropathology characterization in the TgF344-AD rat that reflect preclinical and prodromal AD. Further studies are necessary to investigate more subtle behavioral phenotypes such as detailed sleep architecture. Impulse dyscontrol and agitation are also commonly observed in prodromal AD [35, 50], and can be improved by anti-adernergic drugs [118, 119], which warrants further study in TgF344-AD rats. Changes in behavior in 6-month animals are highly suggestive of altered LC activity in the presence of AD-like hyperphosphorylated tau pathology but in the absence of appreciable forebrain pathology. Current AD therapeutics target late-stage pathology, but targeting prodromal stages with drugs that modulate activity or signaling of early affected structures, such as the LC-NE system, may be more effective at slowing the progression of AD. However, successful development and implementation of these therapies will require a precise understanding of the physiological changes of LC activity across disease stages.

## Acknowledgements

This study was supported by funding from the National Institute of Aging (AG062581 to DW, AG069502 to MAK), the National Institute of Neurological Disorders and Stroke (MS96050 to MAK), and the Eli Lilly Innovation Fellowship Award (CRA and JMR)

The authors of this study would like to thank Ryan McCann and Shivaang Chawla who assisted with rat husbandry.

This study was supported in part by the Rodent Behavioral Core (RBC), which is subsidized by the Emory University School of Medicine and is one of the Emory Integrated Core Facilities. Additional support was provided by the Emory Neuroscience NINDS Core Facilities (P30NS055077). Further support was provided by the Georgia Clinical & Translational Science Alliance of the National Institutes of Health under Award Number UL1TR002378. The content is solely the responsibility of the authors and does not necessarily reflect the official views of the National Institutes of Health.

## Conflict of Interest/Disclosure Statement

The authors have no conflict of interest to report.

## Notes

### Competing Interest Statement

The authors have declared no competing interest.

