## Supplementary Information for "Consequences of endogenous and virally-induced hyperphosphorylated tau on behavior and cognition in a rat model of Alzheimer’s disease"

**Supplemental Information**

| **Primary Antibody** | **Supplier (Product #)** | **Dilution** | **Secondary Antibody** | **Supplier (Product #)** | **Dilution** |
| --- | --- | --- | --- | --- | --- |
| Mouse anti-AT8 | ThermoFisher Scientific (MN1020) | 1:1000 | Goat anti-mouse 488 | ThermoFisher Scientific (A-110001) | 1:500 |
| Mouse anti-4G8 | Biolegend (800709) | 1:1000 | Goat anti-mouse 488 | ThermoFisher Scientific (A-110001) | 1:500 |
| Mouse anti-NET | MAB Technologies (NET05-2) | 1:1000 | Goat anti-mouse 488 | ThermoFisher Scientific (A-110001) | 1:500 |
| Chicken anti-TH | abcam (ab76442) | 1:1000 | Goat anti-chicken 633 | ThermoFisher Scientific (A-21103) | 1:500 |
| Rabbit anti-GFAP | abcam (ab7260) | 1:1000 | Goat anti-rabbit 568 | ThermoFisher Scientific (A-11011) | 1:500 |
| Rabbit anti-IBA1 | FUJIFILM Wako Pure Chemical Corporation (019-19741) | 1:1000 | Goat anti-rabbit 568 | ThermoFisher Scientific (A-11011) | 1:500 |
| Rabbit anti-dsRED | Takara (632496) | 1:1000 | Goat anti-rabbit 568 | ThermoFisher Scientific (A-11011) | 1:500 |

Supplementary Table 1: Antibody Information

Supplementary Table 2: Behavior Statistics

| **Behavior** | **Measure** | **Interaction** | **Main Effect** | **F(DFn, DFd) value** | **p value** |
| --- | --- | --- | --- | --- | --- |
| Sleep Latency | Latency to Fall Asleep | Age x Genotype |  | F (1, 72) = 0.3298 | P=0.5676 |
|  |  | Age x Virus |  | F (1, 72) = 0.2709 | P=0.6043 |
|  |  | Genotype x Virus |  | F (1, 72) = 2.329 | P=0.1313 |
|  |  | Age x Genotype x Virus |  | F (1, 72) = 4.218 | P=0.0436 |
|  |  |  | Age | F (1, 72) = 9.284 | P=0.0032 |
|  |  |  | Genotype | F (1, 72) = 0.2349 | P=0.6294 |
|  |  |  | Virus | F (1, 72) = 0.3153 | P=0.5762 |
| Locomotion | Ambulations, 6 mo - 23 h | Time interval x Genotype |  | F (45, 1665) = 0.8895 | P=0.6807 |
|  |  | Time interval x Virus |  | F (45, 1665) = 0.6977 | P=0.9363 |
|  |  | Genotype x Virus |  | F (1, 37) = 0.08733 | P=0.7692 |
|  |  | Time interval x Genotype x Virus |  | F (45, 1665) = 0.6056 | P=0.9823 |
|  |  |  | Time interval | F (11.84, 437.9) = 31.97 | P<0.0001 |
|  |  |  | Genotype | F (1, 37) = 0.007367 | P=0.9321 |
|  |  |  | Virus | F (1, 37) = 1.019 | P=0.3193 |
|  | Ambulations, 12 mo - 23 h | Time Interval x Genotype |  | F (45, 1575) = 1.118 | P=0.2750 |
|  |  | Time Interval x Virus |  | F (45, 1575) = 1.427 | P=0.0339 |
|  |  | Genotype x Virus |  | F (1, 35) = 0.5669 | P=0.4565 |
|  |  | Time Interval x Genotype x Virus |  | F (45, 1575) = 0.8914 | P=0.6772 |
|  |  |  | Time Interval | F (10.64, 372.3) = 21.61 | P<0.0001 |
|  |  |  | Genotype | F (1, 35) = 0.3077 | P=0.5826 |
|  |  |  | Virus | F (1, 35) = 5.880 | P=0.0206 |
|  | Ambulations, 6 and 12 mo - 23 h | Age x Genotype |  | F (1, 72) = 0.1888 | P=0.6652 |
|  |  | Age x Virus |  | F (1, 72) = 6.304 | P=0.0143 |
|  |  | Genotype x Virus |  | F (1, 72) = 0.6688 | P=0.4162 |
|  |  | Age x Genotype x Virus |  | F (1, 72) = 0.1316 | P=0.7179 |
|  |  |  | Age | F (1, 72) = 4.592 | P=0.0355 |
|  |  |  | Genotype | F (1, 72) = 0.1743 | P=0.6776 |
|  |  |  | Virus | F (1, 72) = 1.336 | P=0.2516 |
|  | Ambulations 6 and 12 mo - Light Phase | Age x Genotype |  | F (1, 72) = 0.2883 | P=0.5930 |
|  |  | Age x Virus |  | F (1, 72) = 0.8304 | P=0.3652 |
|  |  | Genotype x Virus |  | F (1, 72) = 0.3768 | P=0.5412 |
|  |  | Age x Genotype x Virus |  | F (1, 72) = 0.1767 | P=0.6755 |
|  |  |  | Age | F (1, 72) = 0.3634 | P=0.5485 |
|  |  |  | Genotype | F (1, 72) = 0.001673 | P=0.9675 |
|  |  |  | Virus | F (1, 72) = 0.2538 | P=0.6160 |
|  | Ambulations 6 and 12 mo - Dark Phase | Age x Genotype |  | F (1, 72) = 0.4434 | P=0.5076 |
|  |  | Age x Virus |  | F (1, 72) = 4.435 | P=0.0387 |
|  |  | Genotype x Virus |  | F (1, 72) = 0.1508 | P=0.6989 |
|  |  | Age x Genotype x Virus |  | F (1, 72) = 0.2927 | P=0.5902 |
|  |  |  | Age | F (1, 72) = 4.821 | P=0.0313 |
|  |  |  | Genotype | F (1, 72) = 0.7463 | P=0.3905 |
|  |  |  | Virus | F (1, 72) = 0.9755 | P=0.3266 |
|  | Ambulations 6 and 12 mo - Novelty (30 min) | Age x Genotype |  | F (1, 72) = 1.781 | P=0.1862 |
|  |  | Age x Virus |  | F (1, 72) = 3.981 | P=0.0498 |
|  |  | Genotype x Virus |  | F (1, 72) = 0.05367 | P=0.8174 |
|  |  | Age x Genotype x Virus |  | F (1, 72) = 0.7437 | P=0.3913 |
|  |  |  | Age | F (1, 72) = 31.87 | P<0.0001 |
|  |  |  | Genotype | F (1, 72) = 0.5952 | P=0.4429 |
|  |  |  | Virus | F (1, 72) = 2.433 | P=0.1232 |
| Open Field | Perecent Time in Inner Circle | Age x Genotype |  | F (1, 73) = 1.205 | P=0.2759 |
|  |  | Age x Virus |  | F (1, 73) = 3.050 | P=0.0849 |
|  |  | Genotype x Virus |  | F (1, 73) = 0.5330 | P=0.4677 |
|  |  | Age x Genotype x Virus |  | F (1, 73) = 0.2155 | P=0.6439 |
|  |  |  | Age | F (1, 73) = 0.3690 | P=0.5454 |
|  |  |  | Genotype | F (1, 73) = 4.651 | P=0.0343 |
|  |  |  | Virus | F (1, 73) = 3.414 | P=0.0687 |
|  | Total Distance | Age x Genotype |  | F (1, 73) = 0.2590 | P=0.6124 |
|  |  | Age x Virus |  | F (1, 73) = 0.004513 | P=0.9466 |
|  |  | Genotype x Virus |  | F (1, 73) = 0.006385 | P=0.9365 |
|  |  | Age x Genotype x Virus |  | F (1, 73) = 0.05753 | P=0.8111 |
|  |  |  | Age | F (1, 73) = 14.57 | P=0.0003 |
|  |  |  | Genotype | F (1, 73) = 0.4169 | P=0.5205 |
|  |  |  | Virus | F (1, 73) = 0.1072 | P=0.7443 |
| Elevated Plus Maze | Percent Time in Open Arms | Age x Genotype |  | F (1, 70) = 0.2621 | P=0.6103 |
|  |  | Age x Virus |  | F (1, 70) = 0.8899 | P=0.3488 |
|  |  | Genotype x Virus |  | F (1, 70) = 0.001751 | P=0.9667 |
|  |  | Age x Genotype x Virus |  | F (1, 70) = 1.009 | P=0.3186 |
|  |  |  | Age | F (1, 70) = 2.075 | P=0.1542 |
|  |  |  | Genotype | F (1, 70) = 0.6081 | P=0.4381 |
|  |  |  | Virus | F (1, 70) = 1.009 | P=0.3186 |
|  | Total Distance | Age x Genotype |  | F (1, 70) = 1.241 | P=0.2691 |
|  |  | Age x Virus |  | F (1, 70) = 0.02635 | P=0.8715 |
|  |  | Genotype x Virus |  | F (1, 70) = 0.09059 | P=0.7643 |
|  |  | Age x Genotype x Virus |  | F (1, 70) = 0.08218 | P=0.7752 |
|  |  |  | Age | F (1, 70) = 0.08442 | P=0.7723 |
|  |  |  | Genotype | F (1, 70) = 0.6866 | P=0.4101 |
|  |  |  | Virus | F (1, 70) = 0.8085 | P=0.3716 |
| Novelty Suppressed Feeding | Latency in a Novel Cage | Age x Genotype |  | F (1, 73) = 0.1833 | P=0.6698 |
|  |  | Age x Virus |  | F (1, 73) = 0.1495 | P=0.7001 |
|  |  | Genotype x Virus |  | F (1, 73) = 0.04426 | P=0.8340 |
|  |  | Age x Genotype x Virus |  | F (1, 73) = 0.007685 | P=0.9304 |
|  |  |  | Age | F (1, 73) = 9.691 | P=0.0026 |
|  |  |  | Genotype | F (1, 73) = 11.12 | P=0.0013 |
|  |  |  | Virus | F (1, 73) = 0.02327 | P=0.8792 |
|  | Latency in Home Cage | Age x Genotype |  | F (1, 73) = 1.144 | P=0.2883 |
|  |  | Age x Virus |  | F (1, 73) = 6.831e-006 | P=0.9979 |
|  |  | Genotype x Virus |  | F (1, 73) = 1.782 | P=0.1861 |
|  |  | Age x Genotype x Virus |  | F (1, 73) = 0.09143 | P=0.7632 |
|  |  |  | Age | F (1, 73) = 0.1776 | P=0.6747 |
|  |  |  | Genotype | F (1, 73) = 1.454 | P=0.2317 |
|  |  |  | Virus | F (1, 73) = 0.08312 | P=0.7739 |
|  | Pellet Weight Eaten | Age x Genotype |  | F (1, 73) = 1.405 | P=0.2397 |
|  |  | Age x Virus |  | F (1, 73) = 0.8978 | P=0.3465 |
|  |  | Genotype x Virus |  | F (1, 73) = 2.302 | P=0.1335 |
|  |  | Age x Genotype x Virus |  | F (1, 73) = 1.183 | P=0.2803 |
|  |  |  | Age | F (1, 73) = 0.02367 | P=0.8781 |
|  |  |  | Genotype | F (1, 73) = 1.405 | P=0.2397 |
|  |  |  | Virus | F (1, 73) = 0.6447 | P=0.4246 |
| Forced Swim Test | Percent Time Spent Floating | Age x Genotype |  | F (1, 72) = 0.1784 | P=0.6740 |
|  |  | Age x Virus |  | F (1, 72) = 0.004712 | P=0.9455 |
|  |  | Genotype x Virus |  | F (1, 72) = 0.1063 | P=0.7453 |
|  |  | Age x Genotype x Virus |  | F (1, 72) = 0.8174 | P=0.3690 |
|  |  |  | Age | F (1, 72) = 6.904 | P=0.0105 |
|  |  |  | Genotype | F (1, 72) = 0.01323 | P=0.9087 |
|  |  |  | Virus | F (1, 72) = 0.05062 | P=0.8226 |
| Morris Water Maze - Acquisition | Distance to Platform, 6 mo - Training | Day x Genotype |  | F (3, 111) = 2.289 | P=0.0824 |
|  |  | Day x Virus |  | F (3, 111) = 2.188 | P=0.0935 |
|  |  | Genotype x Virus |  | F (1, 37) = 0.8334 | P=0.3672 |
|  |  | Day x Genotype x Virus |  | F (3, 111) = 2.102 | P=0.1040 |
|  |  |  | Day | F (2.759, 102.1) = 44.77 | P<0.0001 |
|  |  |  | Genotype | F (1, 37) = 6.080 | P=0.0184 |
|  |  |  | Virus | F (1, 37) = 0.03508 | P=0.8524 |
|  | Distance to Platform, 12 mo - Training | Day x Genotype |  | F (3, 108) = 1.123 | P=0.3431 |
|  |  | Day x Virus |  | F (3, 108) = 0.3593 | P=0.7825 |
|  |  | Genotype x Virus |  | F (1, 36) = 0.06028 | P=0.8074 |
|  |  | Day x Genotype x Virus |  | F (3, 108) = 0.1379 | P=0.9372 |
|  |  |  | Day | F (2.173, 78.23) = 70.02 | P<0.0001 |
|  |  |  | Genotype | F (1, 36) = 2.709 | P=0.1085 |
|  |  |  | Virus | F (1, 36) = 5.618 | P=0.0233 |
|  | Distance to Platform, 6 and 12 mo - Training | Age x Genotype |  | F (1, 73) = 0.1201 | P=0.7299 |
|  |  | Age x Virus |  | F (1, 73) = 0.3228 | P=0.5717 |
|  |  | Genotype x Virus |  | F (1, 73) = 0.03278 | P=0.8568 |
|  |  | Age x Genotype x Virus |  | F (1, 73) = 0.1442 | P=0.7053 |
|  |  |  | Age | F (1, 73) = 0.05859 | P=0.8094 |
|  |  |  | Genotype | F (1, 73) = 1.556 | P=0.2163 |
|  |  |  | Virus | F (1, 73) = 0.6485 | P=0.4233 |
|  | Distance to Platform, 6 and 12 mo - Probe | Age x Genotype |  | F (1, 73) = 0.07450 | P=0.7857 |
|  |  | Age x Virus |  | F (1, 73) = 0.01041 | P=0.9190 |
|  |  | Genotype x Virus |  | F (1, 73) = 1.727 | P=0.1930 |
|  |  | Age x Genotype x Virus |  | F (1, 73) = 0.02698 | P=0.8700 |
|  |  |  | Age | F (1, 73) = 5.336 | P=0.0237 |
|  |  |  | Genotype | F (1, 73) = 2.193 | P=0.1430 |
|  |  |  | Virus | F (1, 73) = 0.9538 | P=0.3320 |
| Morris Water Maze – Reverse Acquisition | Distance to Platform, 6 mo - Training | Day x Genotype |  | F (3, 111) = 1.592 | P=0.1954 |
|  |  | Day x Virus |  | F (3, 111) = 1.272 | P=0.2875 |
|  |  | Genotype x Virus |  | F (1, 37) = 0.1116 | P=0.7402 |
|  |  | Day x Genotype x Virus |  | F (3, 111) = 1.317 | P=0.2724 |
|  |  |  | Day | F (1.793, 66.33) = 35.49 | P<0.0001 |
|  |  |  | Genotype | F (1, 37) = 4.999 | P=0.0315 |
|  |  |  | Virus | F (1, 37) = 0.01777 | P=0.8947 |
|  | Distance to Platform, 12 mo - Training | Day x Genotype |  | F (3, 108) = 0.5176 | P=0.6711 |
|  |  | Day x Virus |  | F (3, 108) = 1.054 | P=0.3720 |
|  |  | Genotype x Virus |  | F (1, 36) = 0.007100 | P=0.9333 |
|  |  | Day x Genotype x Virus |  | F (3, 108) = 0.4378 | P=0.7264 |
|  |  |  | Day | F (3, 108) = 22.96 | P<0.0001 |
|  |  |  | Genotype | F (1, 36) = 9.539 | P=0.0039 |
|  |  |  | Virus | F (1, 36) = 0.09099 | P=0.7647 |
|  | Distance to Platform, 6 and 12 mo - Training | Age x Genotype |  | F (1, 73) = 0.3419 | P=0.5606 |
|  |  | Age x Virus |  | F (1, 73) = 0.04507 | P=0.8325 |
|  |  | Genotype x Virus |  | F (1, 73) = 0.01787 | P=0.8940 |
|  |  | Age x Genotype x Virus |  | F (1, 73) = 0.002743 | P=0.9584 |
|  |  |  | Age | F (1, 73) = 2.103 | P=0.1513 |
|  |  |  | Genotype | F (1, 73) = 3.363 | P=0.0708 |
|  |  |  | Virus | F (1, 73) = 0.02998 | P=0.8630 |
|  | Distance to Platform, 6 and 12 mo - Probe | Age x Genotype |  | F (1, 72) = 0.3775 | P=0.5409 |
|  |  | Age x Virus |  | F (1, 72) = 0.03185 | P=0.8588 |
|  |  | Genotype x Virus |  | F (1, 72) = 0.1740 | P=0.6778 |
|  |  | Age x Genotype x Virus |  | F (1, 72) = 0.1609 | P=0.6895 |
|  |  |  | Age | F (1, 72) = 0.09493 | P=0.7589 |
|  |  |  | Genotype | F (1, 72) = 0.3182 | P=0.5745 |
|  |  |  | Virus | F (1, 72) = 0.02243 | P=0.8814 |
| Fear Conditioning | Percent Freezing, 6 mo - Training | Interval x Genotype |  | F (6, 222) = 0.08349 | P=0.9978 |
|  |  | Interval x Virus |  | F (6, 222) = 1.189 | P=0.3128 |
|  |  | Genotype x Virus |  | F (1, 37) = 0.8051 | P=0.3754 |
|  |  | Interval x Genotype x Virus |  | F (6, 222) = 0.2719 | P=0.9497 |
|  |  |  | Interval | F (3.948, 146.1) = 43.74 | P<0.0001 |
|  |  |  | Genotype | F (1, 37) = 0.0003098 | P=0.9861 |
|  |  |  | Virus | F (1, 37) = 0.08955 | P=0.7664 |
|  | Percent Freezing, 12 mo - Training | Interval x Genotype |  | F (6, 216) = 2.203 | P=0.0438 |
|  |  | Interval x Virus |  | F (6, 216) = 0.3948 | P=0.8819 |
|  |  | Genotype x Virus |  | F (1, 36) = 0.1900 | P=0.6655 |
|  |  | Interval x Genotype x Virus |  | F (6, 216) = 0.5728 | P=0.7517 |
|  |  |  | Interval | F (3.702, 133.3) = 20.58 | P<0.0001 |
|  |  |  | Genotype | F (1, 36) = 7.184 | P=0.0110 |
|  |  |  | Virus | F (1, 36) = 0.006733 | P=0.9351 |
|  | Percent Freezing, 6 and 12 mo - Training | Age x Genotype |  | F (1, 73) = 0.8330 | P=0.3644 |
|  |  | Age x Virus |  | F (1, 73) = 0.03674 | P=0.8485 |
|  |  | Genotype x Virus |  | F (1, 73) = 0.08266 | P=0.7745 |
|  |  | Age x Genotype x Virus |  | F (1, 73) = 0.4622 | P=0.4987 |
|  |  |  | Age | F (1, 73) = 3.268 | P=0.0748 |
|  |  |  | Genotype | F (1, 73) = 0.9175 | P=0.3413 |
|  |  |  | Virus | F (1, 73) = 0.04438 | P=0.8337 |
|  | Percent Freezing, 6 mo - Context | Interval x Genotype |  | F (6, 222) = 0.9182 | P=0.4826 |
|  |  | Interval x Virus |  | F (6, 222) = 0.3481 | P=0.9105 |
|  |  | Genotype x Virus |  | F (1, 37) = 1.365 | P=0.2502 |
|  |  | Interval x Genotype x Virus |  | F (6, 222) = 0.1483 | P=0.9893 |
|  |  |  | Interval | F (3.852, 142.5) = 10.78 | P<0.0001 |
|  |  |  | Genotype | F (1, 37) = 3.179 | P=0.0828 |
|  |  |  | Virus | F (1, 37) = 0.4996 | P=0.4841 |
|  | Percent Freezing, 12 mo - Context | Interval x Genotype |  | F (6, 216) = 0.4350 | P=0.8550 |
|  |  | Interval x Virus |  | F (6, 216) = 0.6593 | P=0.6826 |
|  |  | Genotype x Virus |  | F (1, 36) = 0.01554 | P=0.9015 |
|  |  | Interval x Genotype x Virus |  | F (6, 216) = 0.3688 | P=0.8982 |
|  |  |  | Interval | F (3.265, 117.6) = 9.797 | P<0.0001 |
|  |  |  | Genotype | F (1, 36) = 0.005031 | P=0.9438 |
|  |  |  | Virus | F (1, 36) = 0.4573 | P=0.5032 |
|  | Percent Freezing, 6 and 12 mo - Context | Age x Genotype |  | F (1, 73) = 0.8824 | P=0.3506 |
|  |  | Age x Virus |  | F (1, 73) = 0.01794 | P=0.8938 |
|  |  | Genotype x Virus |  | F (1, 73) = 0.2401 | P=0.6256 |
|  |  | Age x Genotype x Virus |  | F (1, 73) = 0.4999 | P=0.4818 |
|  |  |  | Age | F (1, 73) = 6.434 | P=0.0133 |
|  |  |  | Genotype | F (1, 73) = 0.5668 | P=0.4540 |
|  |  |  | Virus | F (1, 73) = 0.6819 | P=0.4116 |
|  | Percent Freezing, 6 mo - Cue | Interval x Genotype |  | F (6, 222) = 0.4628 | P=0.8354 |
|  |  | Interval x Virus |  | F (6, 222) = 1.016 | P=0.4158 |
|  |  | Genotype x Virus |  | F (1, 37) = 2.207 | P=0.1459 |
|  |  | Interval x Genotype x Virus |  | F (6, 222) = 1.289 | P=0.2632 |
|  |  |  | Interval | F (3.489, 129.1) = 100.7 | P<0.0001 |
|  |  |  | Genotype | F (1, 37) = 0.5109 | P=0.4792 |
|  |  |  | Virus | F (1, 37) = 0.09123 | P=0.7643 |
|  | Percent Freezing, 12 mo - Cue | Interval x Genotype |  | F (6, 216) = 1.914 | P=0.0797 |
|  |  | Interval x Virus |  | F (6, 216) = 1.723 | P=0.1168 |
|  |  | Genotype x Virus |  | F (1, 36) = 0.4232 | P=0.5195 |
|  |  | Interval x Genotype x Virus |  | F (6, 216) = 0.1969 | P=0.9774 |
|  |  |  | Interval | F (3.348, 120.5) = 49.29 | P<0.0001 |
|  |  |  | Genotype | F (1, 36) = 0.1270 | P=0.7237 |
|  |  |  | Virus | F (1, 36) = 0.7414 | P=0.3949 |
|  | Percent Freezing, 6 and 12 mo - Cue | Age x Genotype |  | F (1, 73) = 0.3804 | P=0.5393 |
|  |  | Age x Virus |  | F (1, 73) = 0.5732 | P=0.4514 |
|  |  | Genotype x Virus |  | F (1, 73) = 1.120 | P=0.2935 |
|  |  | Age x Genotype x Virus |  | F (1, 73) = 0.1419 | P=0.7075 |
|  |  |  | Age | F (1, 73) = 4.431 | P=0.0387 |
|  |  |  | Genotype | F (1, 73) = 0.0001671 | P=0.9897 |
|  |  |  | Virus | F (1, 73) = 0.1547 | P=0.6952 |

Supplementary Table 3: Immunohistochemistry Statistics

| **Stain (Antibody)** | **Region** | **Interaction** | **Main Effect** | **F(DFn, DFd) value** | **p value** |
| --- | --- | --- | --- | --- | --- |
| Hyperphosphorylated Tau (AT8) | LC | Age x Genotype |  | F (1, 34) = 0.1946 | P=0.6619 |
|  |  |  | Age | F (1, 34) = 0.6288 | P=0.4333 |
|  |  |  | Genotype | F (1, 34) = 4.857 | P=0.0344 |
| Amyloid (4G8) | DG | Age x Genotype |  | F (1, 38) = 34.32 | P<0.0001 |
|  |  | Age x Virus |  | F (1, 38) = 1.561 | P=0.2192 |
|  |  | Genotype x Virus |  | F (1, 38) = 0.01764 | P=0.8950 |
|  |  | Age x Genotype x Virus |  | F (1, 38) = 1.665 | P=0.2048 |
|  |  |  | Age | F (1, 38) = 34.36 | P<0.0001 |
|  |  |  | Genotype | F (1, 38) = 92.88 | P<0.0001 |
|  |  |  | Virus | F (1, 38) = 0.03064 | P=0.8620 |
|  | CA3 | Age x Genotype |  | F (1, 38) = 69.32 | P<0.0001 |
|  |  | Age x Virus |  | F (1, 38) = 0.2623 | P=0.6115 |
|  |  | Genotype x Virus |  | F (1, 38) = 0.08914 | P=0.7669 |
|  |  | Age x Genotype x Virus |  | F (1, 38) = 0.2377 | P=0.6287 |
|  |  |  | Age | F (1, 38) = 69.53 | P<0.0001 |
|  |  |  | Genotype | F (1, 38) = 101.1 | P<0.0001 |
|  |  |  | Virus | F (1, 38) = 0.07557 | P=0.7849 |
|  | CA1 | Age x Genotype |  | F (1, 38) = 47.14 | P<0.0001 |
|  |  | Age x Virus |  | F (1, 38) = 0.1195 | P=0.7315 |
|  |  | Genotype x Virus |  | F (1, 38) = 0.5785 | P=0.4516 |
|  |  | Age x Genotype x Virus |  | F (1, 38) = 0.1011 | P=0.7523 |
|  |  |  | Age | F (1, 38) = 46.15 | P<0.0001 |
|  |  |  | Genotype | F (1, 38) = 73.95 | P<0.0001 |
|  |  |  | Virus | F (1, 38) = 0.6275 | P=0.4332 |
| Hyperphosphorylated Tau (AT8) | DG | Age x Genotype |  | F (1, 33) = 2.311 | P=0.1380 |
|  |  | Age x Virus |  | F (1, 33) = 0.1241 | P=0.7269 |
|  |  | Genotype x Virus |  | F (1, 33) = 0.1311 | P=0.7196 |
|  |  | Age x Genotype x Virus |  | F (1, 33) = 0.3596 | P=0.5528 |
|  |  |  | Age | F (1, 33) = 0.2829 | P=0.5983 |
|  |  |  | Genotype | F (1, 33) = 4.467 | P=0.0422 |
|  |  |  | Virus | F (1, 33) = 3.391 | P=0.0746 |
|  | CA3 | Age x Genotype |  | F (1, 38) = 6.305 | P=0.0164 |
|  |  | Age x Virus |  | F (1, 38) = 2.211 | P=0.1453 |
|  |  | Genotype x Virus |  | F (1, 38) = 0.5920 | P=0.4464 |
|  |  | Age x Genotype x Virus |  | F (1, 38) = 0.1392 | P=0.7112 |
|  |  |  | Age | F (1, 38) = 1.593 | P=0.2145 |
|  |  |  | Genotype | F (1, 38) = 2.225 | P=0.1441 |
|  |  |  | Virus | F (1, 38) = 2.137 | P=0.1520 |
|  | CA1 | Age x Genotype |  | F (1, 38) = 7.596 | P=0.0089 |
|  |  | Age x Virus |  | F (1, 38) = 2.817 | P=0.1015 |
|  |  | Genotype x Virus |  | F (1, 38) = 0.7051 | P=0.4063 |
|  |  | Age x Genotype x Virus |  | F (1, 38) = 0.01587 | P=0.9004 |
|  |  |  | Age | F (1, 38) = 8.648 | P=0.0055 |
|  |  |  | Genotype | F (1, 38) = 3.273 | P=0.0783 |
|  |  |  | Virus | F (1, 38) = 0.7262 | P=0.3995 |
| Astrocytes (GFAP) | DG | Age x Genotype |  | F (1, 38) = 12.04 | P=0.0013 |
|  |  | Age x Virus |  | F (1, 38) = 0.1457 | P=0.7048 |
|  |  | Genotype x Virus |  | F (1, 38) = 0.4881 | P=0.4890 |
|  |  | Age x Genotype x Virus |  | F (1, 38) = 0.04013 | P=0.8423 |
|  |  |  | Age | F (1, 38) = 21.56 | P<0.0001 |
|  |  |  | Genotype | F (1, 38) = 36.66 | P<0.0001 |
|  |  |  | Virus | F (1, 38) = 1.334 | P=0.2553 |
|  | CA3 | Age x Genotype |  | F (1, 38) = 14.11 | P=0.0006 |
|  |  | Age x Virus |  | F (1, 38) = 0.2107 | P=0.6488 |
|  |  | Genotype x Virus |  | F (1, 38) = 0.3115 | P=0.5800 |
|  |  | Age x Genotype x Virus |  | F (1, 38) = 0.1110 | P=0.7408 |
|  |  |  | Age | F (1, 38) = 27.91 | P<0.0001 |
|  |  |  | Genotype | F (1, 38) = 31.43 | P<0.0001 |
|  |  |  | Virus | F (1, 38) = 0.2094 | P=0.6499 |
|  | CA1 | Age x Genotype |  | F (1, 38) = 4.730 | P=0.0359 |
|  |  | Age x Virus |  | F (1, 38) = 0.5784 | P=0.4516 |
|  |  | Genotype x Virus |  | F (1, 38) = 0.07893 | P=0.7803 |
|  |  | Age x Genotype x Virus |  | F (1, 38) = 3.977 | P=0.0533 |
|  |  |  | Age | F (1, 38) = 23.76 | P<0.0001 |
|  |  |  | Genotype | F (1, 38) = 5.528 | P=0.0240 |
|  |  |  | Virus | F (1, 38) = 0.2605 | P=0.6127 |
| Microglia (IBA1) | DG | Age x Genotype |  | F (1, 38) = 0.001233 | P=0.9722 |
|  |  | Age x Virus |  | F (1, 38) = 0.09385 | P=0.7610 |
|  |  | Genotype x Virus |  | F (1, 38) = 1.090 | P=0.3030 |
|  |  | Age x Genotype x Virus |  | F (1, 38) = 0.7188 | P=0.4019 |
|  |  |  | Age | F (1, 38) = 0.9495 | P=0.3360 |
|  |  |  | Genotype | F (1, 38) = 8.596 | P=0.0057 |
|  |  |  | Virus | F (1, 38) = 0.01557 | P=0.9014 |
|  | CA3 | Age x Genotype |  | F (1, 39) = 0.8946 | P=0.3501 |
|  |  | Age x Virus |  | F (1, 39) = 0.3941 | P=0.5338 |
|  |  | Genotype x Virus |  | F (1, 39) = 0.2941 | P=0.5907 |
|  |  | Age x Genotype x Virus |  | F (1, 39) = 0.5531 | P=0.4615 |
|  |  |  | Age | F (1, 39) = 3.960 | P=0.0536 |
|  |  |  | Genotype | F (1, 39) = 3.108 | P=0.0857 |
|  |  |  | Virus | F (1, 39) = 3.379 | P=0.0736 |
|  | CA1 | Age x Genotype |  | F (1, 37) = 0.006922 | P=0.9341 |
|  |  | Age x Virus |  | F (1, 37) = 1.323 | P=0.2575 |
|  |  | Genotype x Virus |  | F (1, 37) = 0.5493 | P=0.4633 |
|  |  | Age x Genotype x Virus |  | F (1, 37) = 0.2734 | P=0.6042 |
|  |  |  | Age | F (1, 37) = 24.81 | P<0.0001 |
|  |  |  | Genotype | F (1, 37) = 0.7212 | P=0.4012 |
|  |  |  | Virus | F (1, 37) = 0.9558 | P=0.3346 |
| Norepinephrine Transporter (NET) | DG | Age x Genotype |  | F (1, 38) = 0.05830 | P=0.8105 |
|  |  | Age x Virus |  | F (1, 38) = 0.9959 | P=0.3246 |
|  |  | Genotype x Virus |  | F (1, 38) = 1.277 | P=0.2655 |
|  |  | Age x Genotype x Virus |  | F (1, 38) = 1.814 | P=0.1860 |
|  |  |  | Age | F (1, 38) = 0.01035 | P=0.9195 |
|  |  |  | Genotype | F (1, 38) = 16.36 | P=0.0002 |
|  |  |  | Virus | F (1, 38) = 0.4586 | P=0.5024 |
|  | CA3 | Age x Genotype |  | F (1, 40) = 0.9422 | P=0.3376 |
|  |  | Age x Virus |  | F (1, 40) = 0.6420 | P=0.4277 |
|  |  | Genotype x Virus |  | F (1, 40) = 2.169 | P=0.1487 |
|  |  | Age x Genotype x Virus |  | F (1, 40) = 0.9345 | P=0.3395 |
|  |  |  | Age | F (1, 40) = 8.286 | P=0.0064 |
|  |  |  | Genotype | F (1, 40) = 0.4147 | P=0.5233 |
|  |  |  | Virus | F (1, 40) = 2.742 | P=0.1056 |
|  | CA1 | Age x Genotype |  | F (1, 37) = 1.074 | P=0.3068 |
|  |  | Age x Virus |  | F (1, 37) = 1.183 | P=0.2837 |
|  |  | Genotype x Virus |  | F (1, 37) = 2.548 | P=0.1189 |
|  |  | Age x Genotype x Virus |  | F (1, 37) = 0.4558 | P=0.5038 |
|  |  |  | Age | F (1, 37) = 1.305 | P=0.2607 |
|  |  |  | Genotype | F (1, 37) = 1.846 | P=0.1825 |
|  |  |  | Virus | F (1, 37) = 0.6171 | P=0.4371 |

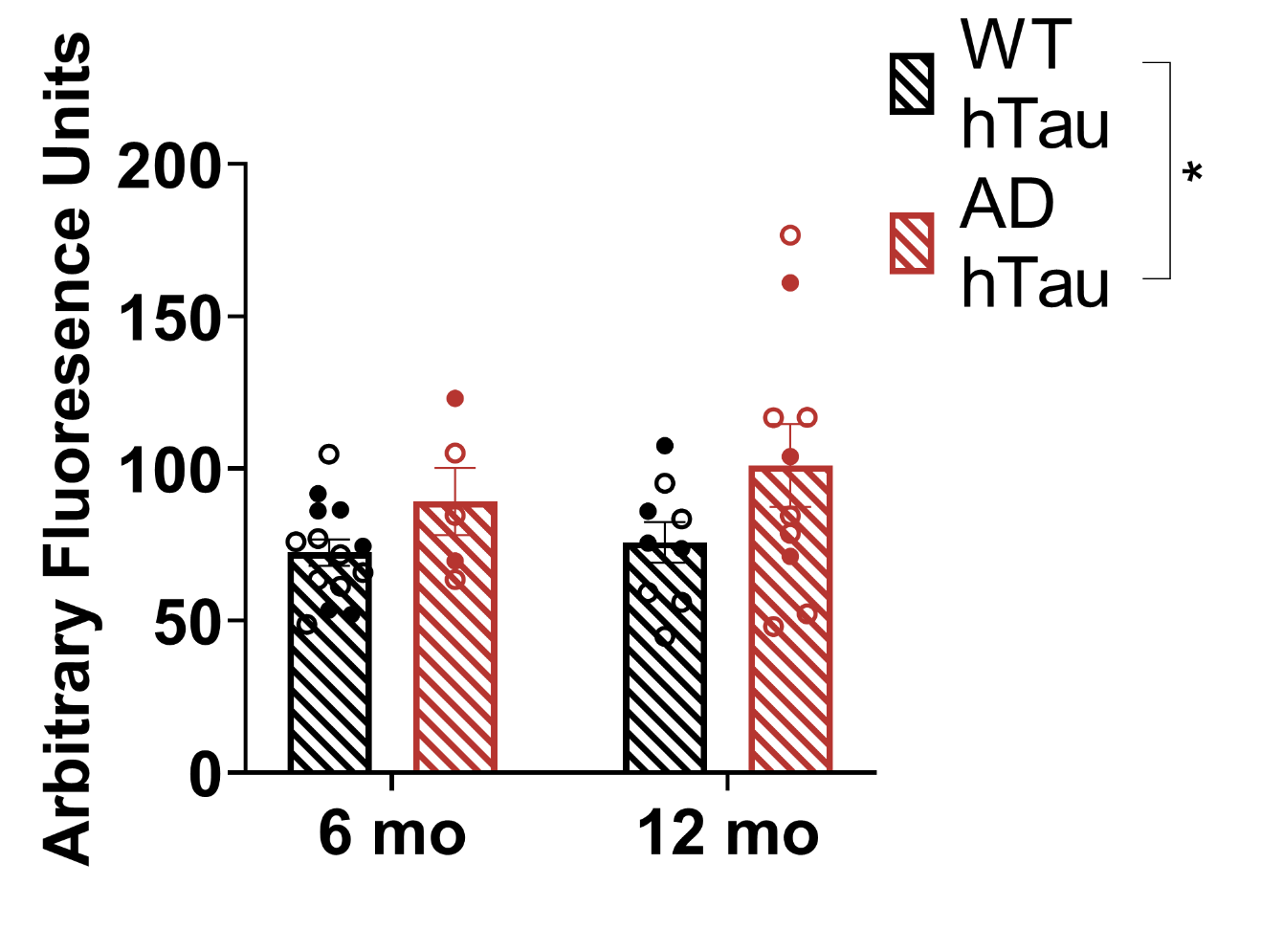

Supplementary Figure 1. Quantification of AT8 pathology in the LC of WT and TgF344-AD littermates.

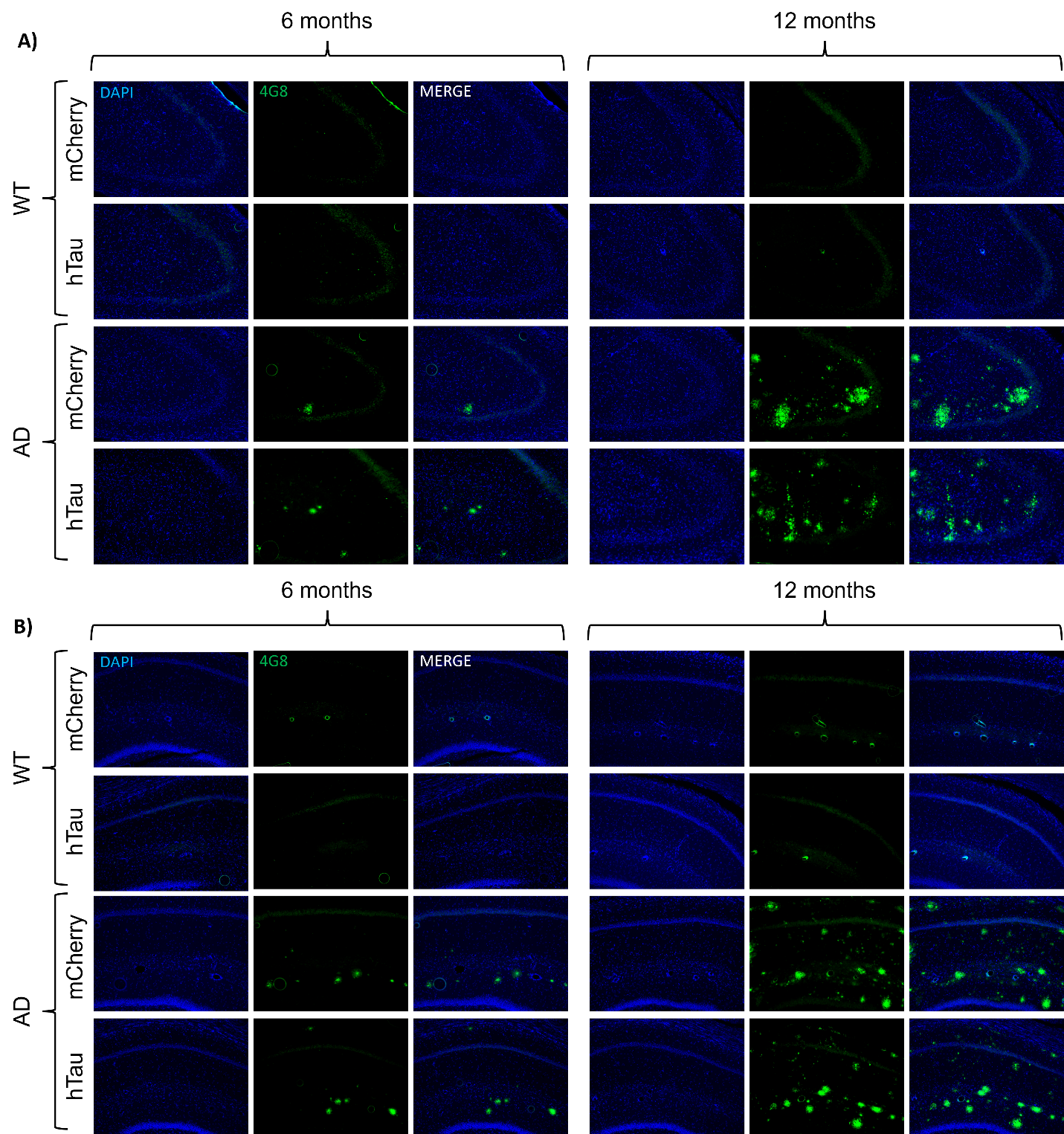

Supplementary Figure 2. Representative images of amyloid pathology in the CA3 (A) and CA1 (B) subregions of the hippocampus. Images taken at 20x magnification.

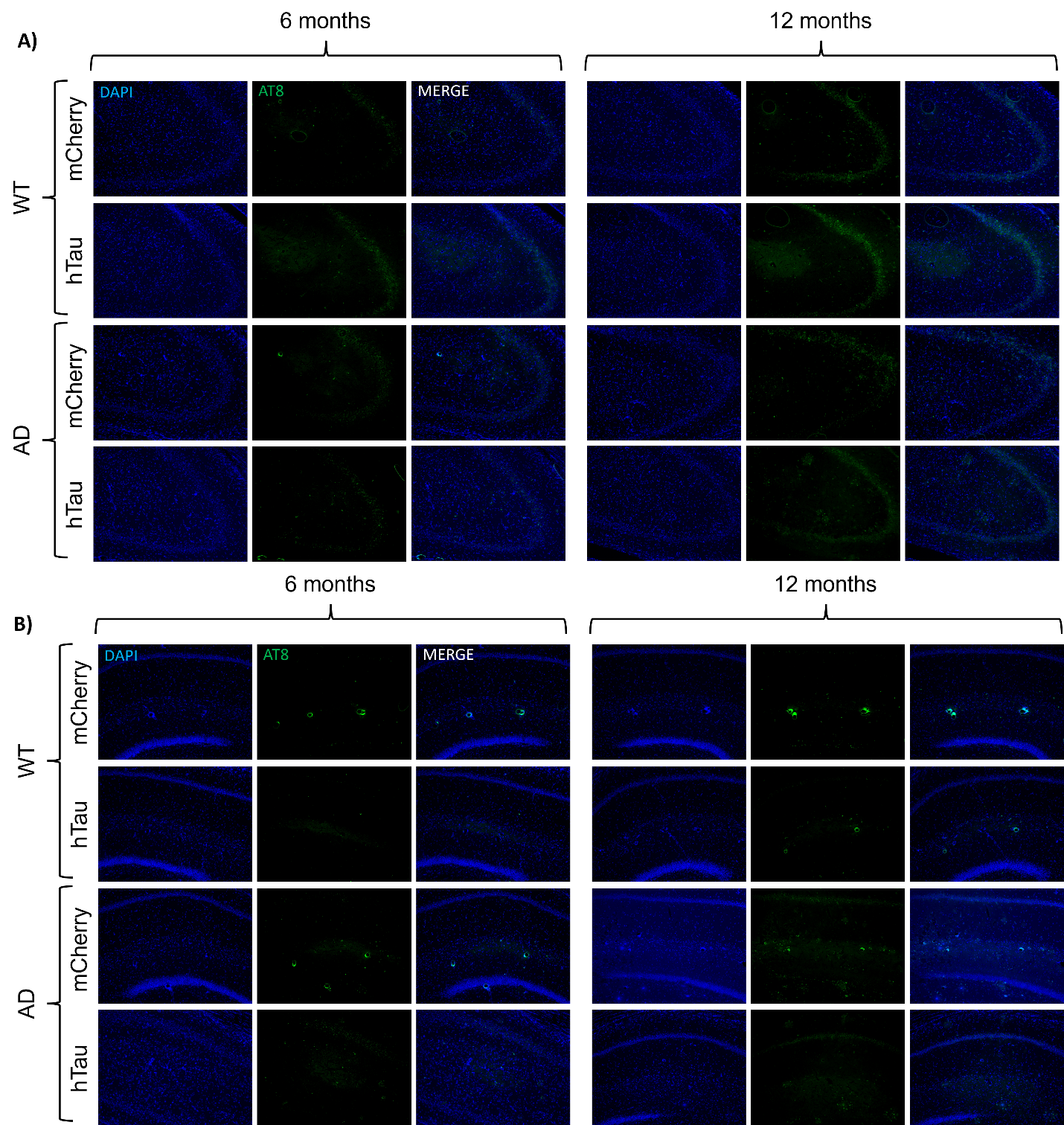

Supplementary Figure 3. Representative images of tau pathology (AT8) in the CA3 (A) and CA1 (B) subregions of the hippocampus. Images taken at 20x magnification.

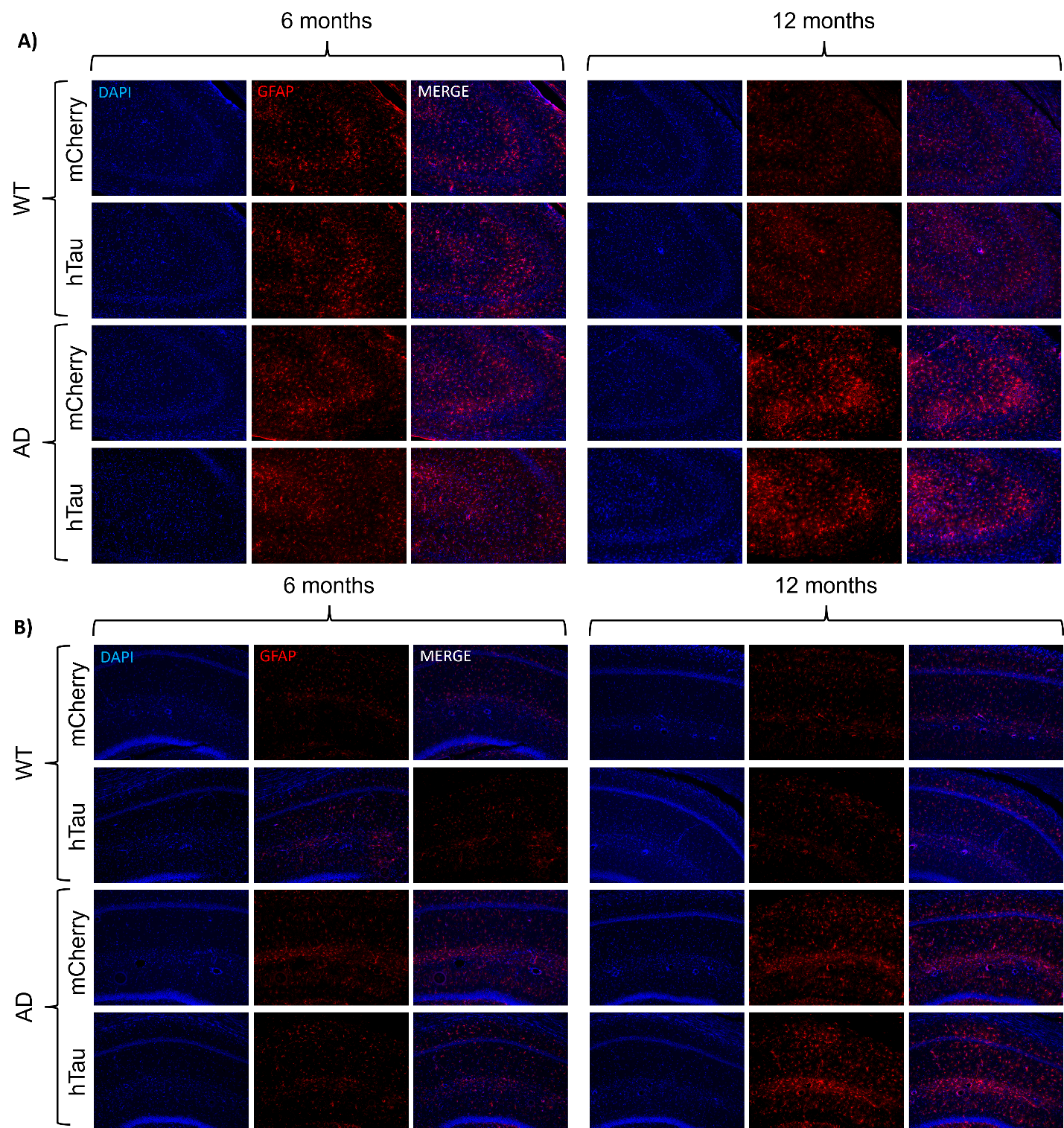

Supplementary Figure 4. Representative images of neuroinflammatory astrocytic marker GFAP in the CA3 (A) and CA1 (B) subregions of the hippocampus. Images taken at 20x magnification.

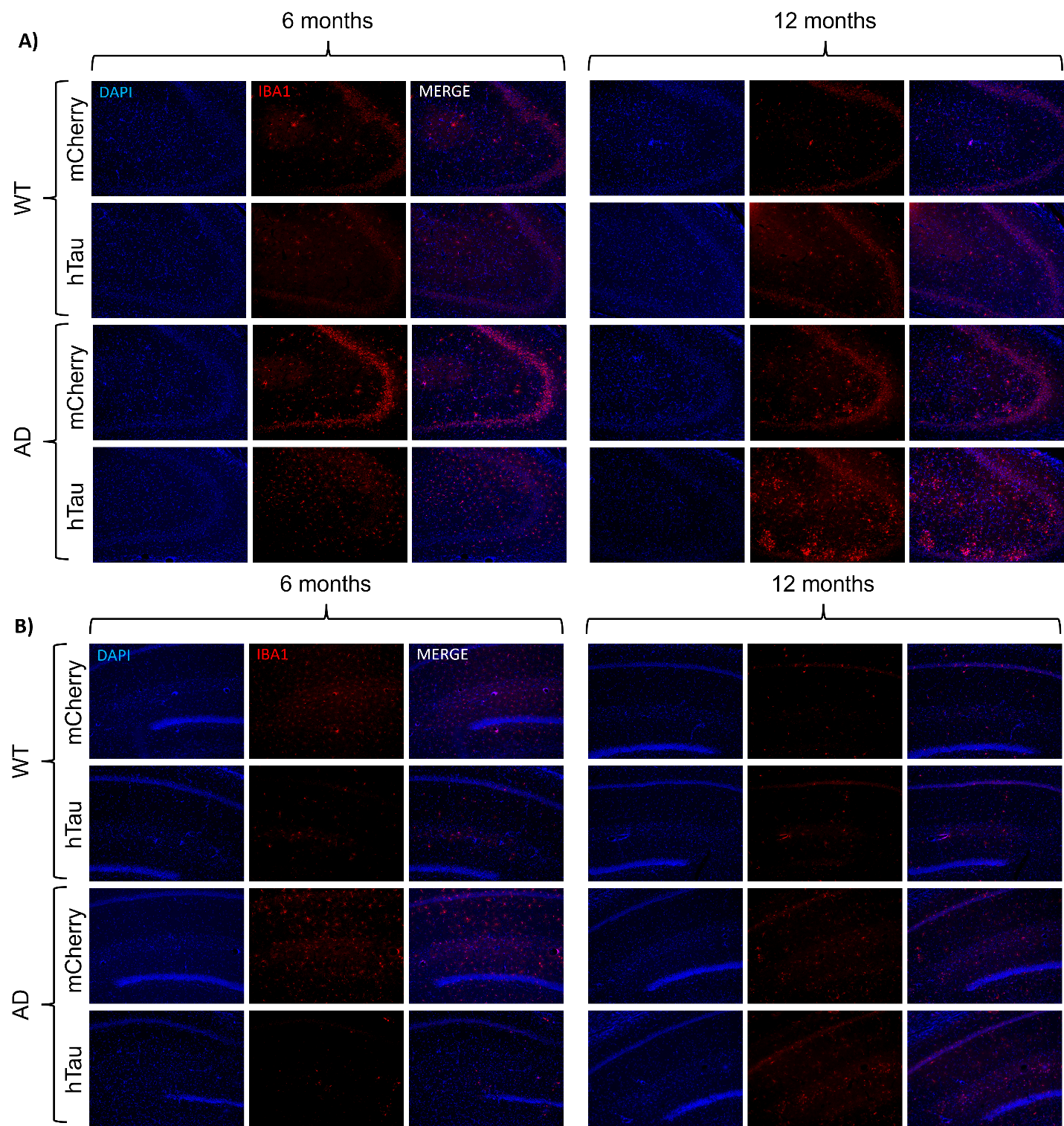

Supplementary Figure 5. Representative images of neuroinflammatory microglial marker IBA1 in the CA3 (A) and CA1 (B) subregions of the hippocampus. Images taken at 20x magnification.

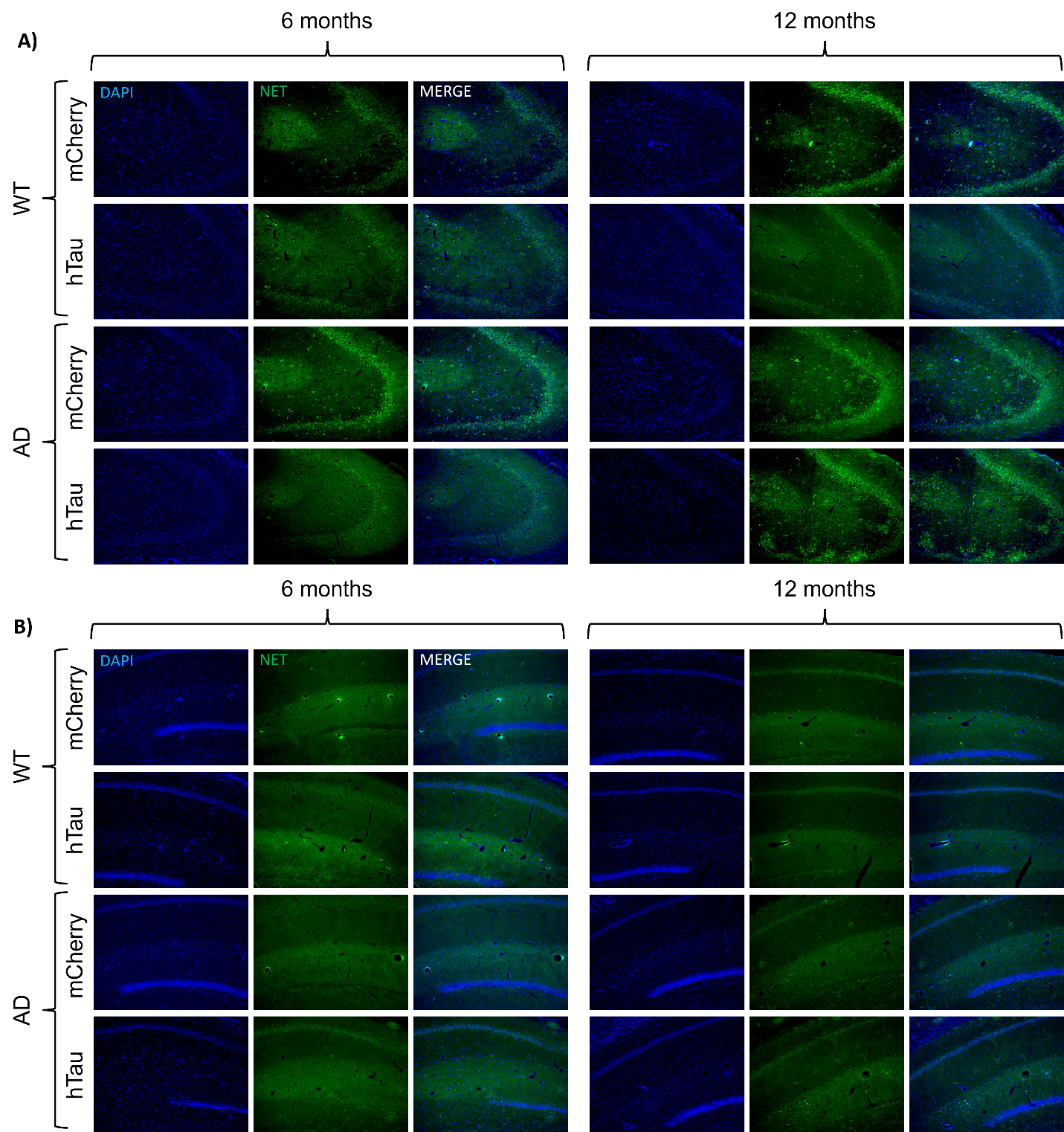

Supplementary Figure 6. Representative images of NET staining in the CA3 (A) and CA1 (B) subregions of the hippocampus. Images taken at 20x magnification.
